# Disruption of the TCA cycle reveals an ATF4-dependent integration of redox and amino acid metabolism

**DOI:** 10.1101/2021.07.27.453996

**Authors:** Dylan G. Ryan, Ming Yang, Hiran A. Prag, Giovanny Rodriguez Blanco, Efterpi Nikitopoulou, Marc Segarra-Mondejar, Christopher A. Powell, Tim Young, Nils Burger, Jan Lj. Miljkovic, Michal Minczuk, Michael P. Murphy, Alexander von Kriegsheim, Christian Frezza

**Affiliations:** MRC Cancer Unit, Hutchison/MRC Research Centre, Cambridge, CB2 0XZ; MRC Mitochondrial Biology Unit, The Keith Peters Building, Cambridge, CB2 0XY; Edinburgh CRUK Centre, Institute of Genetics and Cancer, EH4 2XR; Department of Medicine, University of Cambridge, Cambridge Biomedical Campus, Cambridge, CB2 0QQ

**Keywords:** TCA cycle, reductive carboxylation, fumarate hydratase, succinate dehydrogenase, mitochondria, amino acids, redox homeostasis, respiration, proline, aspartate, asparagine, glutathione, integrated stress response, ATF4

## Abstract

The Tricarboxylic Acid Cycle (TCA) cycle is arguably the most critical metabolic cycle in physiology and exists as an essential interface coordinating cellular metabolism, bioenergetics, and redox homeostasis. Despite decades of research, a comprehensive investigation into the consequences of TCA cycle dysfunction remains elusive. Here, we targeted two TCA cycle enzymes, fumarate hydratase (FH) and succinate dehydrogenase (SDH), and combined metabolomics, transcriptomics, and proteomics analyses to fully appraise the consequences of TCA cycle inhibition (TCAi) in kidney epithelial cells. Our comparative approach shows that TCAi elicits a convergent rewiring of redox and amino acid metabolism dependent on the activation of ATF4 and the integrated stress response (ISR). Furthermore, we also uncover a divergent metabolic response, whereby acute FHi, but not SDHi, can maintain asparagine levels via reductive carboxylation and maintenance of cytosolic aspartate synthesis. Our work highlights an important interplay between the TCA cycle, redox biology and amino acid homeostasis.

**Highlights:**

1. TCA cycle inhibition promotes GSH synthesis and impairs *de novo* aspartate and proline synthesis
2. Disruption of mitochondrial thiol redox homeostasis phenocopies TCA cycle inhibition by promoting GSH synthesis and impairing proline and aspartate synthesis
3. Acute FHi, but not SDHi, can maintain asparagine levels via reductive carboxylation and maintenance of cytosolic aspartate synthesis
4. TCA cycle inhibition mimics an amino acid deprivation-type response and activates ATF4 via the integrated stress response to maintain redox and amino acid homeostasis

## Introduction

Most of the energy-rich adenosine triphosphate (ATP) generated in metabolism is provided by the complete oxidation of key fuel molecules to CO2 in mitochondria. This catabolic process primarily occurs in the mitochondrial matrix via a series of reactions known as the Krebs cycle or the tricarboxylic acid (TCA) cycle (Krebs and Johnson, 1980; Owen et al., 2002; Saraste, 1999). In addition, the TCA cycle is a source of precursors for the synthesis of many other biological molecules, such as nonessential amino acids (NEAAs), lipids, nucleotide bases and porphyrin (Owen *et al*., 2002). Therefore, any dysfunction of the TCA cycle is expected to elicit profound metabolic reprogramming of the cell beyond defective ATP generation. FH is a homotetrameric enzyme localized to the mitochondrial matrix and cytosol that catalyzes the stereospecific hydration of fumarate to L-malate (Tuboi et al., 1986). SDH, also known as complex II of the electron transport chain (ETC), is a heterotetrameric complex composed of four different subunits, SDHA, SDHB, SDHC and SDHD (Yankovskaya et al., 2003). SDH localizes to the inner mitochondrial membrane (IMM) and catalyzes the oxidation of succinate to fumarate (Schultz and Chan, 2001; Yankovskaya *et al*., 2003). SDH is the only TCA cycle enzyme that is also a component of the ETC and so provides a physical link between the TCA cycle and oxidative phosphorylation (OXPHOS). Interestingly, mutations of both FH and SDH predispose kidney tubuluar epithelium to transformation (Sciacovelli et al., 2020), indicating that the kidney can be particularly affected by TCA cycle dysfunction.

Mitochondrial stress activates a retrograde signaling pathway to communicate metabolic dysfunction to the nucleus, a process first described in yeast over twenty years ago (Butow and Avadhani, 2004). However, only recently has mitochondrial stress signalling in mammals been linked to the integrated stress response (ISR) and the transcription factor Atf4 (Bao et al., 2016; Mick et al., 2020; Pakos-Zebrucka et al., 2016; Quirós et al., 2017). The ISR is an evolutionary conserved signaling network that responds to diverse cellular stresses, from amino acid deprivation to viral infection, and operates by reprogramming translation (general translation inhibitor via phosphorylation of the translational initiation factor eIF2α) and increasing the translation of specific stress-induced mRNAs, such as *Atf4* (Pakos-Zebrucka *et al*., 2016). While much is understood about the mechanisms of ISR signalling and Atf4 activation as a whole, we are only beginning to understand how mitochondria engage this pathway and how the ISR and Atf4 regulate the metabolic response to stress. Likewise, despite decades of research, a comprehensive investigation into the metabolic consequences and cellular response to TCA cycle dysfunction specifically in mammalian cells remains elusive. Here, we perform a comparative investigation into the consequences of acute TCAi in murine epithelial kidney cells. We identify a shared metabolic signature of TCAi, whereby enhanced glutathione synthesis is accompanied by a concomitant impairment in de novo proline and aspartate synthesis. Importantly, the selective disruption of mitochondrial thiol redox homeostasis is sufficient to recapitulate the convergent TCAi metabolic signature. Furthermore, acute TCAi phenocopied an amino acid deprivation-like state and activated the integrated stress response to counter redox and amino acid stress. Notably, acute SDHi led to a pronounced decrease in asparagine synthesis, an effect not observed with FHi, where reductive carboxylation and cytosolic aspartate synthesis maintain the asparagine pool. Finally, this work highlights a previously underappreciated role for oxidative TCA cycle activity and respiration in proline synthesis.

## Results

### TCA cycle inhibition promotes glutathione synthesis while impairing de novo proline and aspartate synthesis

To identify a common metabolic signature of TCA cycle dysfunction, we performed liquid chromatography-mass spectrometry (LC-MS)-based metabolomic analysis of kidney epithelial cells upon TCAi. To this aim, we assessed the response to acute TCAi with the FH inhibitor, FHIN-1, and the SDH inhibitors, Atpenin A5 (AA5) and Thenoyltrifluoroacetone (TTFA) (Acute model) (Figure 1A) as it enabled us to chart the early events of TCAi in a controlled fashion without adaptive compensations. We then compared it to cells in which *Fh1* or *Sdhb* have been genetically ablated (Chronic model). This analysis revealed substantial changes in the metabolome upon TCAi (Figure S1A–S1H) with *bona fide* metabolic markers of FH and SDH deficiency observed (Figure S1E-H). For FHi, this included the accumulation of (S)-2-succinocysteine (2SC) and succinicGSH (Figure S1E and G), metabolites formed from fumarate-mediated alkylation of free cysteine and the major anti-oxidant GSH (Zheng et al., 2015). For SDHi, this included a significant accumulation in intracellular succinate levels (Figure S1F and H). Comparative analysis of all five conditions revealed that only two metabolites were significantly increased (Figure 1B), whilst eight metabolites were significantly decreased across all conditions (Figure 1C). The two significantly increased metabolites were GSH and oxidized GSH (GSSG), which suggested a perturbation in redox homeostasis when TCA cycle activity is inhibited. Of the eight decreased metabolites, notably, two were TCA cycle metabolites (malate and *cis*-aconitate). Furthermore, the amino acids proline and aspartate were also decreased. Completing the list were propionylcarnitine and butyrylcarnitine, which are products of catabolic pathways known to feed the TCA cycle, while UTP and UDP-glucose are pyrimidine-associated metabolites (synthesized from aspartate) involved in glucose metabolism. Interestingly, while both acute FHi and SDHi impaired intracellular aspartate, proline and cystine levels (albeit to a different extent), only SDHi led to a significant decrease in intracellular asparagine levels (Figure 1G-H and Figure S1I). Of note, acute FHi or SDHi led to a decrease in aspartate and proline levels already after one hour of treatment (Figure S1J and K), which suggests that the impairment in amino acid synthesis precedes the changes in GSH metabolism. Finally, we observed that *Fh1*^-/-^ and *Sdhb*^-/-^ cells increased intracellular cystine, serine, glycine, and asparagine levels, suggesting that chronic perturbations of the TCA cycle may induce adaptive compensatory metabolic changes (Figure 1D-E).

**Figure 1.**
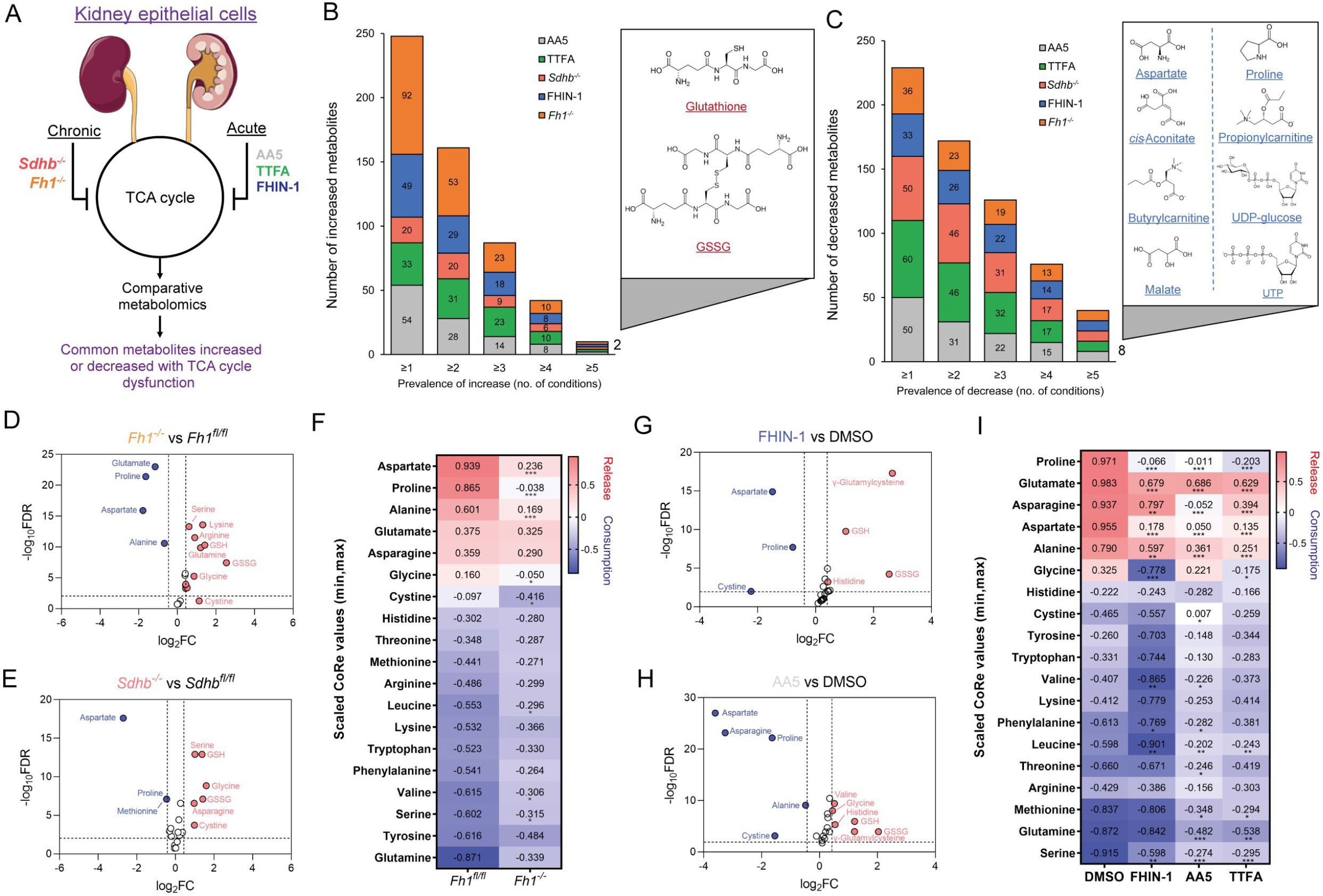
TCA cycle inhibition promotes cytosolic glutathione synthesis while impairing de novo proline and aspartate synthesis. (A) Schematic diagram of comparative metabolomic approach to assess the metabolic response of kidney epithelial cells to TCAi. (B) Significantly increased metabolites (Cut-off = 25% change in abundance, false discovery rate (FDR) = 5%) with genetic or pharmacological TCAi. (C) Significantly decreased metabolites. (B-C) (24 h timepoint). (D) Volcano plot of glutathione-related metabolites and amino acids in *Fh1*^-/-^ versus *Fh1^fl/fl^* cells. (E) Volcano plot of glutathione-related metabolites and amino acids in *Sdhb*^-/-^ versus *Sdhb^fl/fl^* cells. (F) Heatmap of mean scaled consumption-release (CoRe) intensity value of amino acids in *Fh1*^-/-^ versus *Fh1^fl/fl^*cells. (B-F) (*n* = 5-10 independent biological replicates). (G) Volcano plot of glutathione-related metabolites and amino acids in FHIN-1 (20 μM)-treated *Fh1^fl/fl^* versus DMSO-vehicle control *Fh1^fl/fl^* cells. (H) Volcano plot of glutathione-related metabolites and amino acids in AA5 (1 μM)-treated *Fh1^fl/fl^* versus DMSO-treated *Fh1^fl/fl^* cells. (G-H) (24 h timepoint) (*n* = 10-15 independent biological replicates). (I) Heatmap of mean scaled CoRe intensity value of amino acids in FHIN-1, AA5 and TTFA (500 μM) versus DMSO-treated *Fh1^fl/fl^* cells (24 h timepoint) (*n* = 5 independent biological replicates). (D-I) p value determined by multiple unpaired t-tests, corrected with two-stage step-up method of Benjamini, Krieger and Yekutieli - FDR = 5%. p <0.05*; p <0.01**; p < 0.001***.

Considering the profound effect of TCAi on intracellular amino acid levels, we performed a consumption/release (CoRe) experiment upon acute FHi and SDHi. This analysis demonstrated a significant decrease in the release of aspartate, alanine, glutamate and proline (Figure 1I) upon TCAi. Of note, while there was a small but significant decrease in asparagine release upon FHi, SDHi led to a far more pronounced decrease (Figure 1I). Coupled to the intracellular metabolomics, this result confirms a divergent metabolic response to the acute TCAi on asparagine synthesis depending on the enzyme targeted. Consistent with acute TCAi, *Fh1*^-/-^ cells also demonstrated significant impairment in the release of aspartate, alanine and proline (Figure 1F). Overall, our metabolomic analyses identified perturbed redox homeostasis and amino acid metabolism, most notably a diminished capacity to synthesize aspartate and proline and an increase in GSH synthesis, as shared metabolic features of TCAi.

### TCA cycle inhibition and impaired mitochondrial thiol redox homeostasis rewire glutamine metabolism

To further characterize the metabolic reprogramming events associated with acute TCAi, we employed stable-assisted isotope tracing of U-^13^C-glutamine with LC-MS analysis of our kidney epithelial cells, and focused on the three metabolic modules highlighted above, ie proline, aspartate, and GSH metabolism (Figure 2A).

**Figure 2.**
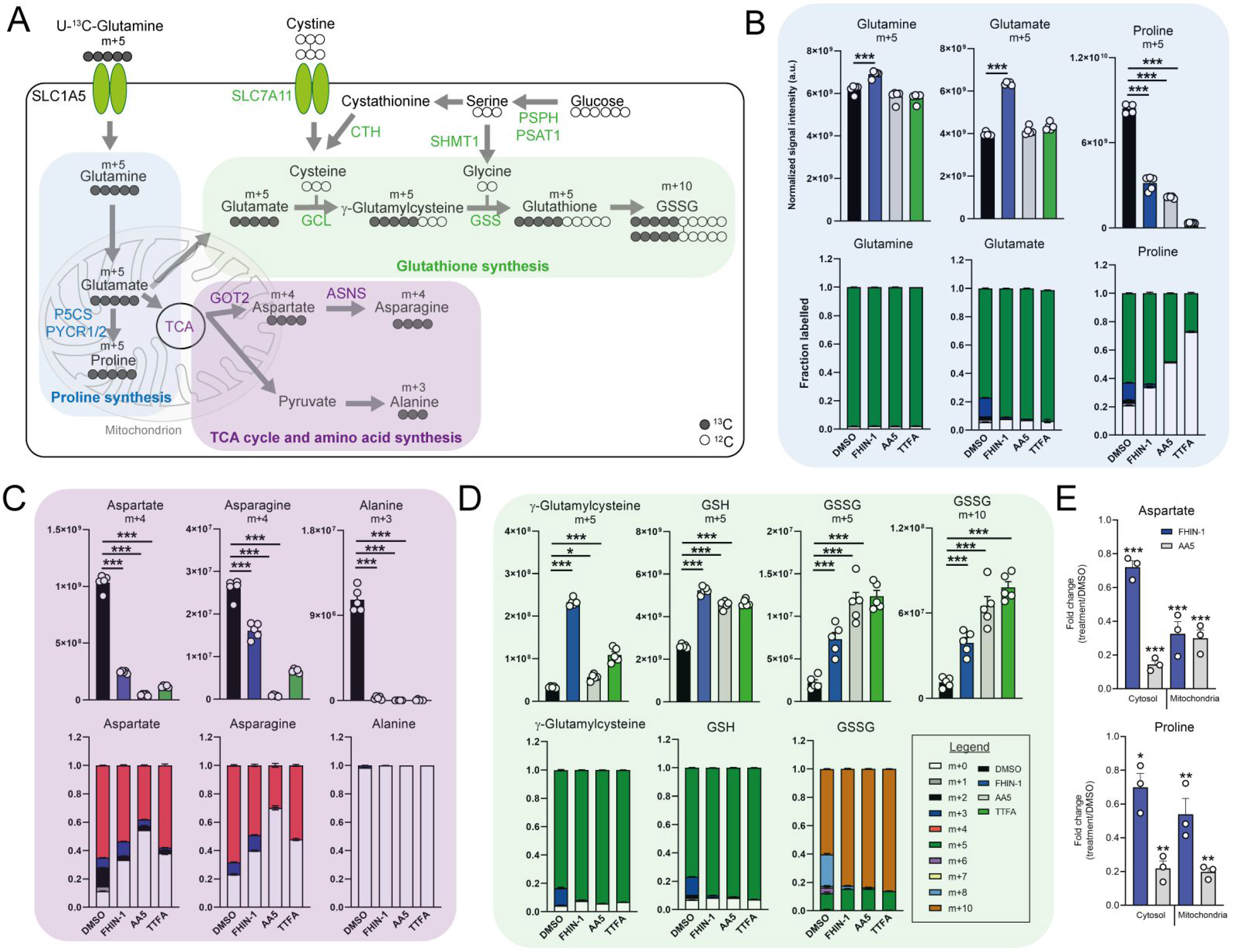
TCA cycle inhibition impairs glutamine-derived proline and aspartate synthesis but promotes cytosolic GSH synthesis. (A) Schematic diagram highlighting U-^13^C-glutamine tracing into distinct metabolic modules. (B) U-^13^C-glutamine tracing into proline (m+5 labelling intensity and total isotopologue fraction distribution). (C) U-^13^C-glutamine tracing into aspartate, asparagine and alanine (m+4 and m+3 labelling intensity and total isotopologue fraction distribution). (D) U-^13^C-glutamine tracing into glutathione synthesis pathway (m+5 and m+10 labelling intensity and total isotopologue fraction distribution) (B-D) show DMSO, FHIN-1 (20 μM)-, AA5 (1 μM)- and TTFA (500 μM)-treated *Fh1^fl/fl^* cells (24 h timepoint) (*n* = 5 independent biological replicates). Data are mean ± standard error of mean (SEM). p value determined by ordinary one-way ANOVA, corrected for multiple comparisons using Tukey statistical hypothesis testing. (E) Proline and aspartate level fold change in mitochondrial and cytosol fractions in FHIN-1 and AA5-treated versus DMSO-treated *Fh1^fl/fl^* cells (24 h timepoint) (*n* = 3 independent biological replicates). Data are mean ± standard error of mean (SEM). p values determined by unpaired two-tailed t-test. p <0.05*; p <0.01**; p < 0.001***.

Glutamine can be converted into glutamate by glutaminase and can then be diverted into several metabolic pathways, including GSH synthesis, proline synthesis, and the TCA cycle (Figure 2A). Stable-assisted isotope tracing revealed that a significant proportion of glutamate and proline are derived from glutamine as their pools consist primarily of the m+5 isotopologue (Figure 2B). Total m+5 labelling from glutamine significantly decreased in proline, suggesting an impairment in glutamate-derived proline synthesis upon TCAi (Figure 2B). Notably, m+3 labelled proline, which is derived from α-ketoglutarate (α-KG) that has undergone one round of oxidative TCA cycle catabolism (Figure S2A), disappears with TCAi. U-^13^C-glutamine tracing also revealed robust labelling of TCA cycle intermediates (Figure S2B) and an impairment in oxidative TCA cycling with FHi and SDHi, as determined by a decrease in m+4 and m+2-labelled citrate and *cis*-aconitate, m+3-labelled α-KG and 2-hydroxyglutarate (2-HG) and m+2-labelled succinate, malate, fumarate and aspartate (Figure S2B). Aspartate is directly synthesized from TCA cycle-derived oxaloacetate via glutamate-aspartate transaminase 2 (GOT2), downstream of FH and SDH in mitochondria (Birsoy et al., 2015; Cardaci et al., 2015; Sullivan et al., 2015). Aspartate is subsequently exported to the cytosol where it can also be converted into asparagine via the action of asparagine synthetase (ASNS) (Zhang et al., 2014). We observed a significant decrease in m+4-labelled aspartate and asparagine upon TCAi, suggesting an impairment in mitochondrial-synthesized aspartate (Figure 2C). It is worth noting that SDHi was more effective at decreasing m+4-labelled aspartate than FHi, explaining the lower levels of asparagine synthesis observed with SDHi. Acute FHi promoted reductive carboxylation, as determined by an increase in m+5-labelled isotopologue fractional enrichment in α-KG, 2-HG, citrate and *cis*-aconitate and in m+3-labelled malate, aspartate and asparagine (which arises from the cytosolic synthesis of aspartate via GOT1) (Birsoy *et al*., 2015) (Figure 2C and S2B). Despite a reduction in the m+3 labelling, the total pools of malate, aspartate, and asparagine were maintained (Figure S2C). In contrast to FHi, SDHi significantly decreased reductive carboxylation and fully impaired cytosolic aspartate and asparagine synthesis (Figure S2C). A complete loss of m+3 labelled alanine was also observed with TCAi, which may be derived from oxaloacetate-derived phosphoenolpyruvate (PEP) or from m+3-labelled malate as a consequence of reductive carboxylation (Figure S2A). Consistent with elevated levels of cytosolic GSH synthesis following TCAi, an increase in the intensity of m+5 labelling in γ-glutamylcysteine, GSH and GSSG and m+10-labelled GSSG was observed (Figure 2D).

This tracing analysis was further supported by combining metabolomics of mitochondrial and cytosolic fractions. Indeed, whilst mitochondrial aspartate levels where equally reduced by both FHi and SDHi, SDHi led to a more pronounced decrease in the cytosolic fraction of aspartate (Figure 2E). Additionally, asparagine levels were maintained in the cytosol with FHi, but decreased with SDHi (Figure S2D). Therefore, asparagine levels are maintained upon FHi in part from a less severe depletion of cytosolic aspartate levels and from the maintenance of reductive carboxylation and GOT1-derived aspartate synthesis. Fractionation also revealed a decrease in proline upon TCAi, and an increase in GSSG levels with SDHi and in succinicGSH with FHi across both fractions (Figure 2E and S2D).

Due to increased oxidative stress with TCAi, as determined by a significant increase in GSSG levels when compared to GSH levels (Figure S3A) and the aforementioned accumulation of succinicGSH adducts with FHi, we investigated whether cytosolic GSH synthesis and amino acid metabolism were sensitive to perturbations in mitochondrial thiol redox homeostasis. To achieve this, we utilized a recently developed tool, termed mitoCDNB, to selectively deplete mitochondrial GSH pools and inhibit thioredoxin reductase 2 and peroxiredoxin 3, thus impairing mitochondrial thiol redox homeostasis (Booty et al., 2019; Cvetko et al., 2020). We confirmed the previously-described formation of the mitoCDNB-GSH adduct (MitoGSDNB) upon treatment with mitoCDNB (Booty *et al*., 2019), thus validating the approach (Figure 3A). Similar to TCAi, impaired mitochondrial thiol redox homeostasis significantly decreased proline and aspartate levels and increased GSH synthesis (Figure 3B). Furthermore, mitoCDNB decreased m+5 labelling in proline and m+4 labelling in aspartate while increasing m+5 labelling in γ-glutamylcysteine, GSH and GSSG and m+10-labelled GSSG (Figure 3C) upon incubation with U-^13^C-glutamine. The decrease in aspartate with mitoCDNB is likely due to reduced oxidative TCA cycling (Figure 3C and S3B). Interestingly, m+4 labelled asparagine was unaffected with mitoCDNB and similarly to FHi, we observe an increase in reductive carboxylation, cytosolic aspartate synthesis, and an increase in m+3-labelled asparagine despite lower levels of m+3-labelled aspartate (Figure S3C). Overall, this data supports the idea that impaired mitochondrial redox homeostasis may causally contribute to the changes in amino acid metabolism observed upon TCAi.

**Figure 3.**
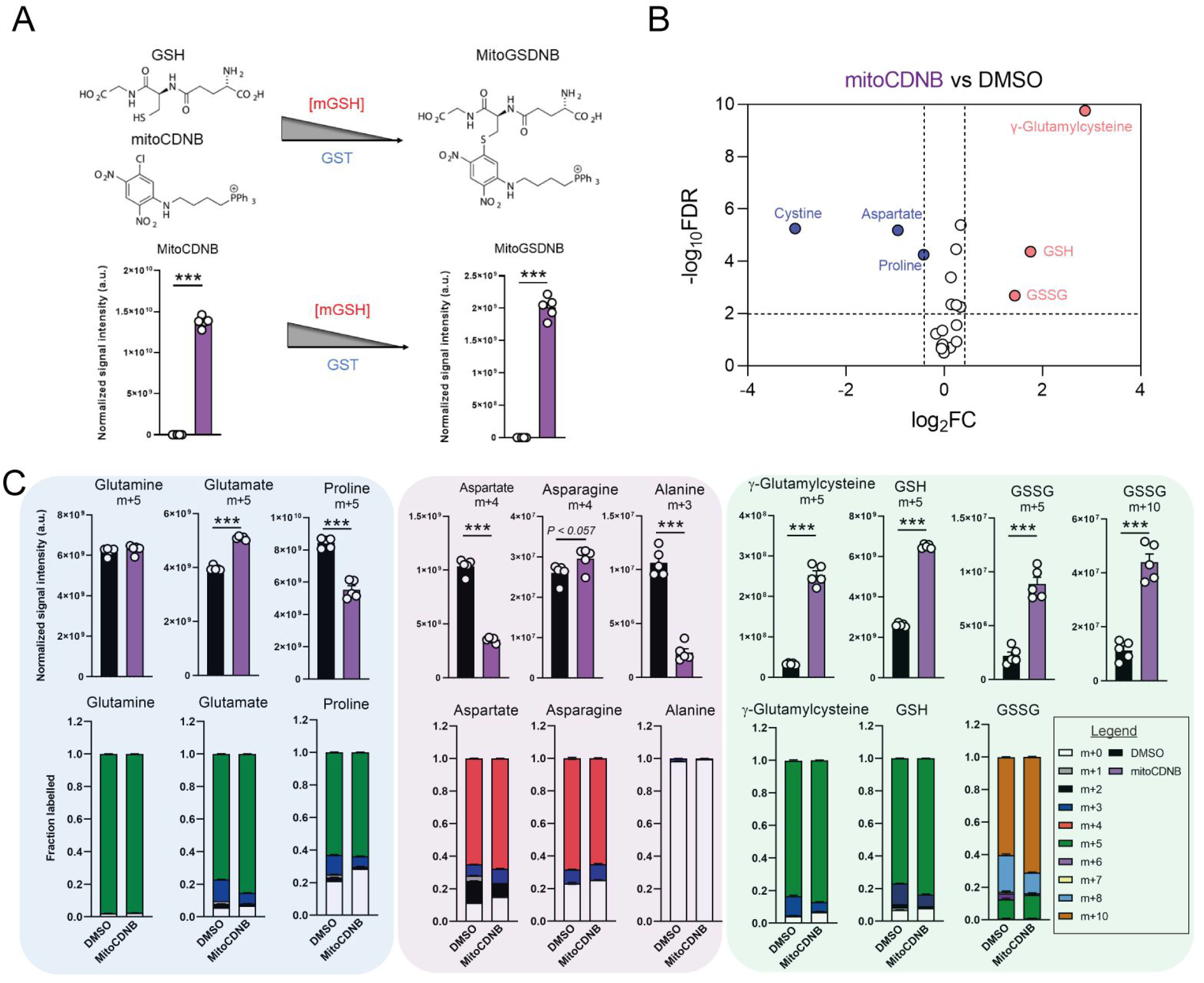
Disrupted mitochondrial thiol redox homeostasis phenocopies TCA cycle inhibition. (A) Schematic diagram highlighting mito 1-Chloro-2,4-dinitrobenzene (mitoCDNB) reaction with glutathione and the formation of mito 1-S-glutathionyl-2,4-dinitrobenzene (MitoGSDNB) and intensity levels in DMSO-treated and mitoCDNB (10 μM)-treated *Fh1*^fl/fl^ cells. (B) Volcano plot of glutathione-related metabolites and amino acids in mitoCDNB-treated *Fh1^fl/fl^* versus DMSO-treated *Fh1^fl/fl^* cells. p value determined by multiple unpaired t-tests, corrected with two-stage step-up method of Benjamini, Krieger and Yekutieli - FDR = 5%. (C) U-^13^C-glutamine tracing (m+10, m+5, m+4 and m+3 labelling intensity and total isotopologue fraction distribution) in DMSO and mitoCDNB-treated *Fh1^fl/fl^* cells. (A-C) (24 h timepoint) (*n* = 5-10 independent biological replicates). (A and C) Data are mean ± SEM. p value determined by unpaired two tailed t-test or ordinary one-way ANOVA corrected for multiple comparisons using Tukey statistical hypothesis testing. p <0.05*; p <0.01**; p < 0.001***.

### Impaired respiration underlies defect in proline biosynthesis

We then investigated the metabolic determinants of the unexpected defects in proline metabolism elicited by TCAi. Given that glutamate is an essential source for proline biosynthesis, TCA cycle anaplerosis, and GSH synthesis, we hypothesized that the defect in proline might arise from differences in glutamate apportioning into these pathways (Figure 2A). We therefore tested whether the *γ*-glutamylcysteine ligase (GCL) inhibitor L-buthionine-sulfoximine (BSO), which blunted GSH synthesis (Figure 4A), would restore proline levels. In support of the glutamate apportioning hypothesis, BSO treatment of kidney epithelial cells increased glutamate, aspartate and asparagine abundance, whilst a trend in increased proline was also observed (Figure 4B). In contrast, when TCAi was coupled with impaired GSH synthesis, it led to significantly higher intracellular glutamate and cystine levels but failed to rescue the decrease in proline (Figure 4C and D). In fact, proline levels decreased further, suggesting that glutathione synthesis is required to support proline synthesis when TCA cycle activity is interrupted and redox stress increases. Likewise, alanine, asparagine and aspartate levels were also significantly lower with a combination of FHi and BSO (Figure 4C), whilst SDHi and BSO lead to a further drop in asparagine (Figure 4D). Similar to BSO treatment, the incubation with ethylGSH (eGSH), which supplements the GSH pool and spares glutathione-related metabolites, such as *γ*-glutamylcysteine, cystine and cys-gly (Figure S4A), increased intracellular glutamate levels when combined with TCAi but failed to rescue the decrease in proline (Figure S4B). Together, these findings show that intracellular glutamate apportioning does not explain the defect in de novo proline synthesis with TCAi.

**Figure 4.**
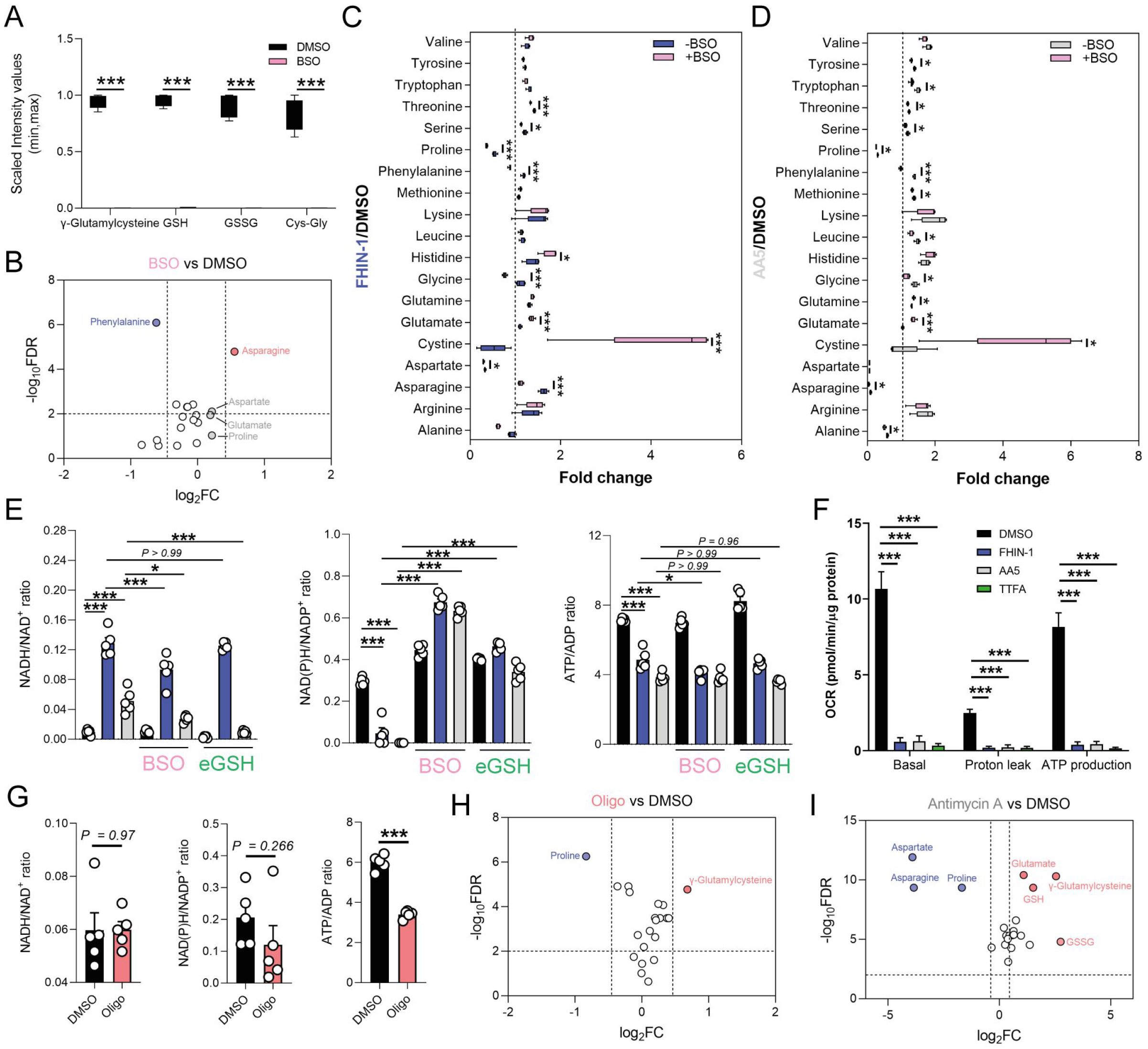
Impaired mitochondrial respiration underlies defect in de novo proline synthesis. (A) Scaled intensity values of glutathione-related metabolites in BSO (500 μM)-treated and DMSO-treated *Fh1^fl/fl^* cells. (B) Volcano plot of amino acids in BSO-treated versus DMSO-treated *Fh1^fl/fl^* cells. (C) Interleaved box and whiskers plot of amino acids with FHIN-1-treated or FHIN-1 and BSO co-treatment of *Fh1^fl/fl^* cells. (D) Interleaved box and whiskers plot of amino acids with AA5-treated or AA5 and BSO cotreatment of *Fh1^fl/fl^* cells. (E) NADH/NAD^+^, NAD(P)H/NADP^+^ and ATP/ADP ratio in DMSO, FHIN-1- and AA5-treated *Fh1^fl/fl^* cells ± BSO or ethylGSH co-treatment. (F) OCR assessed by Seahorse analyser in DMSO, FHIN-1-, AA5- and TTFA-treated *Fh1^fl/fl^* cells (A-F) (24 h timepoint) (*n* = 5 independent biological replicates). (E-F) Data are mean ± SEM. p value determined by ordinary one-way ANOVA, corrected for multiple comparisons using Tukey statistical hypothesis testing. p <0.05*; p <0.01**; p < 0.001***. (G) NADH/NAD^+^, NAD(P)H/NADP^+^ and ATP/ADP ratio in DMSO and Oligomycin (10 μM)-treated *Fh1^fl/fl^* cells. Data are mean ± SEM. p values determined by unpaired two-tailed t-test. p <0.05*; p <0.01**; p < 0.001***. (H) Volcano plot of glutathione-related metabolites and amino acids in Oligomycin-treated versus DMSO-treated *Fh1^fl/fl^* cells (G-H) (1 h timepoint) (*n* = 5 independent biological replicates). (I) Volcano plot of glutathione-related metabolites and amino acids in Antimycin A (2 μM)-treated versus DMSO-treated *Fh1^fl/fl^* cells (24 h timepoint) (*n* = 5 independent biological replicates). (A-D; H-I) p value determined by multiple unpaired t-tests, corrected with two-stage step-up method of Benjamini, Krieger and Yekutieli - FDR = 5%.

To further understand what could give rise to the observed defects in proline synthesis with TCAi, we decided to investigate changes in both the redox and bioenergetic state of the cell. To achieve this, we measured the NADH/NAD^+^, NAD(P)H/NADP^+^ and ATP/ADP ratio with TCAi on its own or coupled to BSO or eGSH treatment (Figure 4E). Intriguingly, we observed a significant increase in the NADH/NAD^+^ ratio with both FHi (~12-fold) and SDHi (~4.65-fold), but the increase was more than double with FHi (Figure 4E). This difference was also reflected by the elevated lactate/pyruvate ratio observed with FHi compared to SDHi, whereas the decrease in the citrate/pyruvate ratio was similar between both conditions (Figure S4D). Furthermore, the NADH/NAD^+^ ratio was decreased with BSO co-treatment suggesting an essential interplay between GSH synthesis and the redox state of the cell upon TCAi (Figure 4E). TCAi also led to a significant decrease in the NAD(P)H/NADP^+^ ratio (Figure 4E). The NAD(P)H/NADP^+^ ratio is a readout of oxidative stress given the role of NAD(P)H in supporting anti-oxidant defence systems and in the reduction of cystine to cysteine for GSH synthesis (Zheng *et al*., 2015). Indeed, we observed an increase in the NAD(P)H/NADP^+^ ratio (and an increase in cystine) with TCAi and BSO co-treatment, while eGSH supplementation prevented a decrease in the NAD(P)H/NADP^+^ ratio with TCAi (Figure 4E). This data suggests that TCAi places cells under conditions of oxidative stress and that the drop in NAD(P)H is due in part to an increased requirement of the cell to convert oxidized cystine to reduced cysteine to support GSH synthesis. Recently, NAD(P)H has been implicated as the major cofactor supporting mitochondrial proline biosynthesis and this could partly explain the decrease in proline (Tran et al., 2021; Zhu et al., 2021). However, neither BSO or eGSH could rescue proline synthesis despite increasing intracellular glutamate or re-establishing NAD(P)H levels (Figure 4B and C). Whilst not precluding the NAD(P)H/NADP^+^ ratio as being an important mitochondrial factor governing this response, it suggested other mechanisms could also be at play with TCAi.

The main function of the TCA cycle is to support NADH generation for OXPHOS. As expected, TCAi led to a significant decrease in the ATP/ADP ratio (Figure 4E) and a decrease in mitochondrial respiration and ATP synthesis, as assessed with a Seahorse flux analyzer (Figure 4F). Unlike the NAD(P)H/NADP^+^ ratio, BSO and eGSH treatments failed to rescue the decrease in the ATP/ADP ratio, and FHi and BSO treatment actually led to a further decrease (Figure 4E). Given the importance of ATP as a cofactor for the proline synthetic enzyme Aldh18a1 (also known as P5C synthetase), this data suggested that impaired respiration and ATP availability was a factor governing reduced proline biosynthesis with TCAi. To validate this hypothesis, we treated cells with the F_1_F_0_-ATP synthase inhibitor oligomycin. In line with our hypothesis, we observed a significant decrease in the ATP/ADP ratio (Figure 4G) and proline (Figure 4H). We also observed an increase in *γ*-glutamylcysteine (Figure 4H) and no significant change in the NADH/NAD^+^ or NAD(P)H/NADP^+^ ratio (Figure 4G), although a trend towards a decrease in the NAD(P)H/NADP^+^ ratio was observed. Given the requirement of ATP for NADK2-mediated phosphorylation of NAD^+^ to generate NADP^+^ in mitochondria (Tran *et al*., 2021; Zhu *et al*., 2021), it’s likely that a combined reduction of mitochondrial ATP and NAD(P)H give rise to the defect in proline synthesis. Further support for mitochondrial respiration involvement in regulating de novo proline synthesis came from the inhibition of complex III with antimycin A, which significantly decreased aspartate, asparagine and proline, while promoting GSH synthesis (Figure 4I). Overall, TCAi uncovered an underappreciated interconnection between GSH and proline metabolism, underpinned by both metabolic and bioenergetics cues.

### TCA cycle inhibition mimics an amino acid deprivation-like response and promotes compensatory reprogramming

We then intended to investigate whether any transcriptional or translational response contributed to the metabolic changes we described upon TCAi. To achieve this, we performed mRNA-seq on FHIN-1- and AA5-treated cells (Figure 5A and C and S5A) and proteomics analysis on FHIN-1- and TTFA-treated cells (Figure 5C and S5B and C). Both treatments resulted in highly significant changes in the transcriptome and gene set enrichment analysis (GSEA) (Subramanian et al., 2005) identified robust activation of an amino acid deprivation response, in line with the decrease in aspartate and proline we observed. We also revealed a heme regulated inhibitor (HRI) stress response signature upon TCAi (Figure 5A). We validated several of the most significantly increased targets via qPCR (Figure 5B) and confirmed that TTFA similarly increased the targets when compared with AA5 (Figure 5B). To identify a common transcriptional and proteomic signature of TCAi, we compared the significantly increased and decreased genes/proteins and constructed comparative Venn diagrams (Figure 5C and S5A and B). Overrepresentation analysis (ORA) of the significantly increased genes/proteins using Enrichr (Chen et al., 2013) revealed that cytosolic tRNA aminoacylation and amino acid metabolism were among the most enriched pathways using the Reactome database (Figure 5C) again lending support to the idea that perturbed amino acid metabolism was a key outcome of interrupted TCA cycle activity. We also observed a suppression of cholesterol biosynthesis (Figure S5A) consistent with previous reports using respiration inhibitors (Mick *et al*., 2020), while mitoribosome subunits and respiration-associated proteins were found to decrease in abundance with TCAi (Figure S5B). However, despite this apparent decrease in mitoribosome subunits, we failed to observe a decrease in 35S-methionine labelling of mitochondrial proteins with TCAi (Figure S5D), which suggests that mitochondrial protein synthesis is resistant to both impaired respiration and a decrease in the components of the mitochondrial translation apparatus over the tested timeframe (24 h). Notably, FHi led to a more pronounced decrease in Complex I biogenesis and subunits (Figure S5C) consistent with the previous reports on FH-deficiency regulating iron-sulphur cluster biogenesis and complex I activity (Tong et al., 2011; Tyrakis et al., 2017), while also explaining the noticeable differences in the NADH/NAD^+^ ratio between FHi and SDHi (Figure 4E).

**Figure 5.**
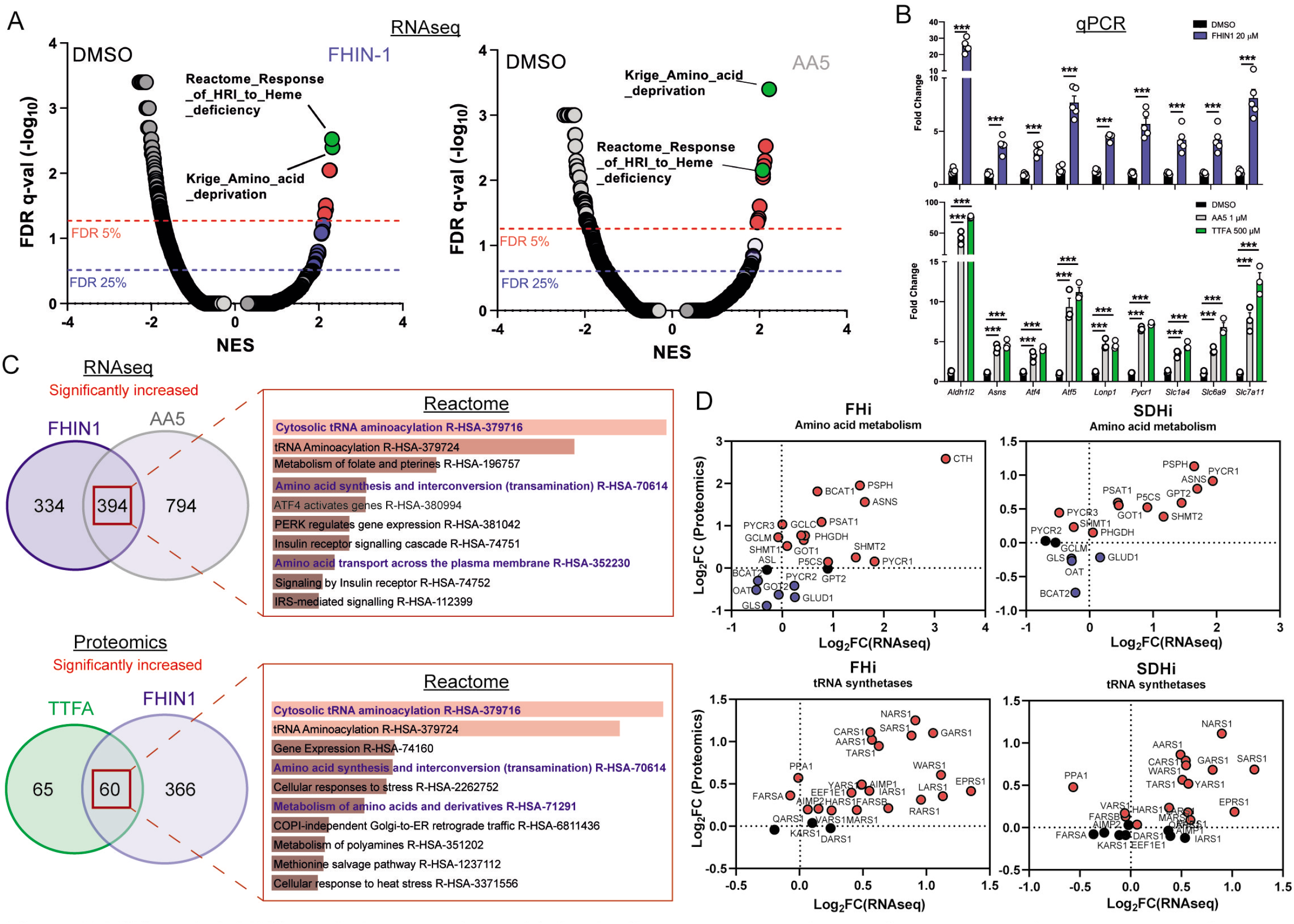
TCA cycle inhibition mimics an amino acid deprivation response and promotes compensatory reprogramming. (A) GSEA of RNAseq data from FHIN-1- and AA5-treated versus DMSO-treated *Fh1^fl/fl^* cells. (B) Quantitative PCR results from FHIN-1-, AA5-, TTFA-, and DMSO-treated *Fh1^fl/fl^* cells (24 h timepoint) (*n* = 3-5 independent biological replicates). Data are mean ± SEM. p value determined by unpaired two-tailed t-test or ordinary one-way ANOVA, corrected for multiple comparisons using Tukey statistical hypothesis testing. p <0.05*; p <0.01**; p < 0.001***. (C) ORA of overlapping transcripts significantly increased in FHIN-1 and AA5-treated *Fh1^fl/fl^* cells and between overlapping proteins significantly increased in FHIN-1 and TTFA-treated *Fh1^fl/fl^* cells (24 h timepoint) (*n* = 3-5 independent biological replicates). Reactome pathways ranked by p value. (D) Comparison of amino acid metabolism and tRNA synthetase transcript and protein levels.

Comparing the log2 fold change of enzymes involved in amino acid metabolism detected in both the proteomics and RNAseq highlighted an increase in the transcript and protein level of Asns with both FHi and SDHi (Figure 5D). Likewise, GOT1 levels were also increased (Figure 5D). This data suggested that cytosolic asparagine synthesis was maintained with FHi in part via reductive carboxylation and GOT1-directed aspartate synthesis, but also due to increased Asns expression. It also emphasized the importance of cytosolic aspartate synthesis, as acute SDHi robustly increased Asns but failed to maintain asparagine synthesis. Interestingly, TCAi led to a robust increase in the cytosolic proline synthetic enzyme, Pycr3, at the protein level. Consistent with a more pronounced decrease in proline levels with SDHi (Figure 1G and H, S1I), Aldh18a1 and Pycr1 were also upregulated to a greater extent at the protein level (Figure 5D). Furthermore, the most highly increased tRNA synthetase at the protein level with TCAi was the asparaginyl-tRNA synthetase (NARS1), while one of the most highly increased on the transcript level was the glutamate-proline tRNA synthetase (EPRS1). These data support the notion that a key metabolic module affected by TCA cycle dysfunction is amino acid metabolism, with a particular focus on proline and aspartate/asparagine. Finally, a notable increase in the cysteine (CARS)-, glycine (GARS)-, alanine (AARS)- and serine(SARS)-tRNA synthetases along with enzymes associated with cysteine, glycine, serine and alanine biosynthesis (CTH, SHMT1/2, PSAT1, PSPH1 and GPT2 with either FHi or SDHi (or both) (Figure 5D) lend further support to the idea TCAi promotes compensatory transcriptional/translational rewiring to support GSH and amino acid metabolism in order to counteract amino acid stress.

### TCA cycle inhibition activates Atf4 and the integrated stress response

To understand how the transcriptional changes induced upon TCAi were communicated to the nucleus, we performed transcription factor (TF) enrichment analysis and uncovered activation of several metabolic stress response TFs associated with the ISR, namely Ddit3/Chop, Atf5, Atf4, Atf3, Cebpb and Cebpg (Figure 6A and B). Using a mitochondrial ISR/Atf4 gene signature curated from the literature (Bao *et al*., 2016; Guo et al., 2020; Quirós *et al*., 2017), we confirmed a significant enrichment of Atf4 targets with both FHi (Figure S6A) and SDHi (Figure S6B) when compared to other hallmark genesets from the molecular signatures database (MsigDB) (Subramanian *et al*., 2005). To validate engagement of the ISR with TCAi, we probed for puromycinylated proteins and phosphorylation of eIF2α on serine 51, a readout of global translation rates and ISR activation (Rabouw et al., 2019), and demonstrated a time-dependent decrease in translation and increase in p-eIF2α as early as 1 h post-treatment (Figure 6C). Furthermore, a robust and time-dependent increase in Atf4 stabilization was also observed (Figure 6C). In addition, mitochondrial GSH depletion with mitoCDNB also activated Atf4 to a similar extent as TCAi (Figure S6C).

**Figure 6.**
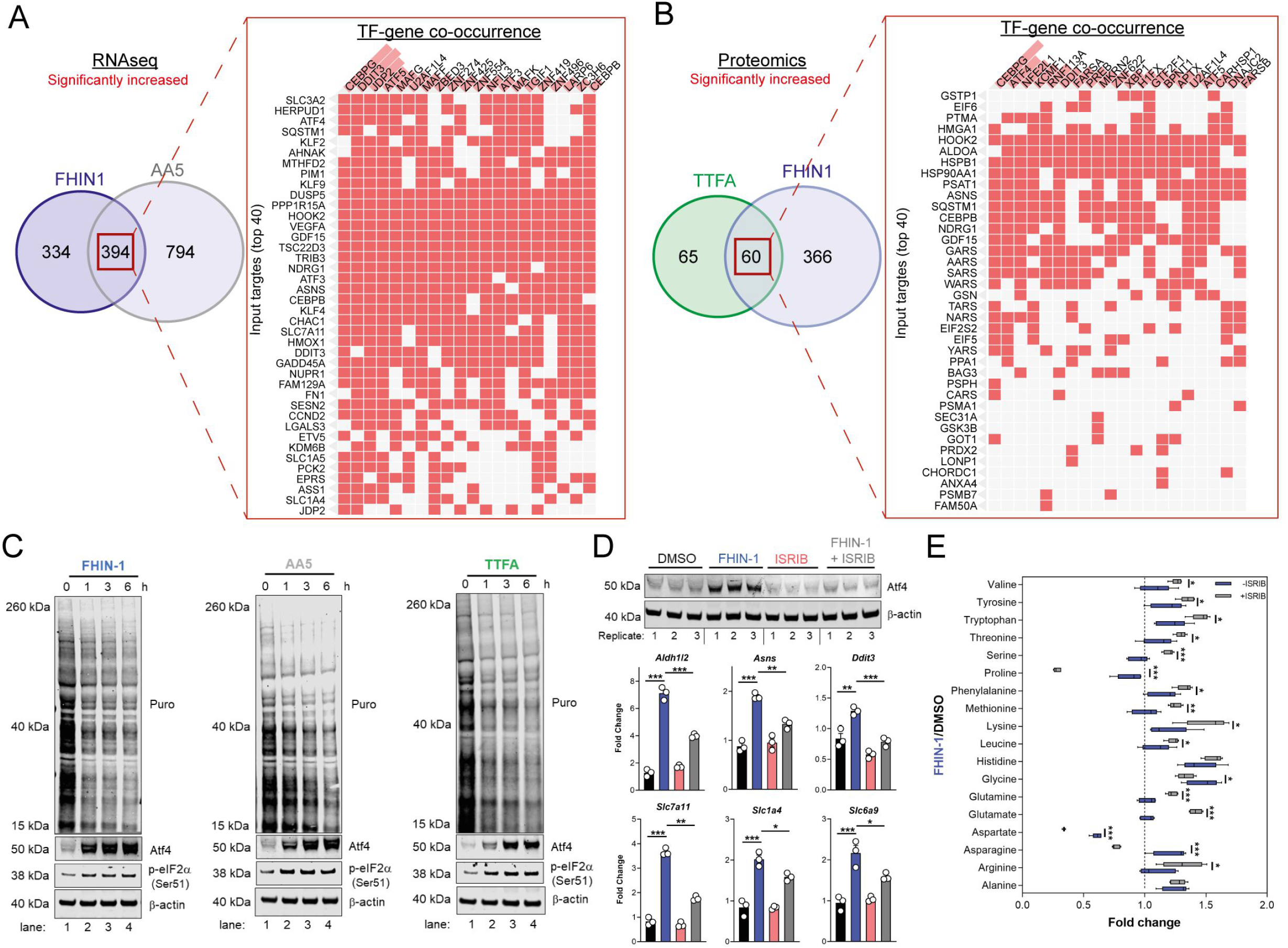
TCA inhibition activates the integrated stress response to counter amino acid and redox stress. (A) Clustergram showing Transcription-factor (TF)-gene co-occurrence ORA of overlapping transcripts significantly increased in FHIN-1 and AA5-treated *Fh1^fl/fl^* cells. (B) Clustergram showing TF-gene co-occurrence ORA of overlapping transcripts significantly increased in FHIN-1 and TTFA-treated *Fh1*^fl/fl^ cells. (A-B) (24 h timepoint) (*n* = 3-5 independent biological replicates). TFs are ranked by combined score (p value and Z score). (C) Western blot analysis of puromycin incorporation, Atf4, phopho-eIF2α and β-actin as loading control in FHIN-1, AA5 and TTFA timecourse treatment of *Fh1^fl/fl^* cells (representative image of 3 independent biological replicates). (D) Western blot analysis of Atf4 and β-actin as loading control (6 h timepoint) and qPCR analysis of Atf4-target genes (24 h timepoint) (*n* = 3 independent biological replicates) in DMSO-, ISRIB (500 nM)-, FHIN-1 (10 μM) and FHIN-1+ISRIB treated *Fh1^fl/fl^* cells. Data are mean ± SEM. p value determined by ordinary one-way ANOVA, corrected for multiple comparisons using Tukey statistical hypothesis testing. (E) Interleaved box and whiskers plot of amino acids with FHIN-1-treated or FHIN-1- and ISRIB-co-treated *Fh1^fl/fl^* cells (24 h timepoint) (*n* = 5 independent biological replicates). p value determined by multiple unpaired t-tests, corrected with two-stage step-up method of Benjamini, Krieger and Yekutieli - FDR = 5%. p <0.05*; p <0.01**; p < 0.001***.

To confirm a role for the ISR in regulating Atf4 stabilization and the expression of Atf4-targets with TCAi, we co-treated cells with FHIN-1 and the specific integrated stress response inhibitor (ISRIB) (Sidrauski et al., 2015), a potent antagonist of p-eIF2α. ISRIB works within a defined window of activation, inhibiting low level ISR activity but not affecting strong ISR signalling (Rabouw *et al*., 2019). Therefore, we used a lower dose of FHIN-1 (10 μM) and observed an attenuation in FHi-induced Atf4 stability and Atf4-target gene expression (Figure 6D). Importantly, this regulation was only strongly observed with FHi-induced Atf4 and not basal Atf4 levels (Figure 6D). This result confirmed that the ISR was involved in Atf4 stabilization and FHi-induced gene expression. In order to determine whether induction of the ISR regulated the metabolic response to TCAi, we treated wildtype cells with FHIN-1 or FHIN-1 and ISRIB (Figure 6E). While ISRIB treatment led to a significant decrease in proline (~25%) (Figure S6D) likely due to basal p-eIF2α (Figure 6C). Co-treatment of FHIN-1 and ISRIB led to a substantial decrease in intracellular proline (~75%) when compared to FHIN-1 alone (Figure 6E). Similarly, aspartate and asparagine levels were also decreased with co-treatment of FHIN-1 and ISRIB (Figure 6E) supporting the idea that the ISR was activated to compensate for the defect in the de novo synthesis of these amino acids. In addition to countering amino acid stress, we observed an increase in oxidative stress with FHi when the ISR was attenuated, as assessed by a significant increase in GSSG/GSH ratio (Figure S6E). In summary, TCAi triggers the ISR to counter amino acid and redox stress in kidney epithelial cells.

To evaluate the role of Atf4 in the response to TCAi, we ablated Atf4 expression using two independent siRNAs and confirmed a decrease at the protein and transcript level of Atf4 and Atf4-target genes (Figure 7A). Atf4 was stabilized to a certain extent basally in our cells and its ablation revealed that Atf4 is a key regulator of GSH synthesis, as expected given it’s important role in regulating cysteine metabolism (Sbodio et al., 2018; Torrence et al., 2021), and of the amino acids proline, alanine and asparagine (Figure 7B). Interestingly, aspartate levels increased with Atf4 knockdown in keeping with the role of Atf4 in upregulating *Asns* expression (Figure 7A). At the same time, intracellular glutamine levels increased in line with an impairment in GSH synthesis (Figure 7B). A comparison of the fold change in amino acids with TCAi, with or without Atf4 silencing, revealed that maintenance of asparagine synthesis was dependent on Atf4-induced *Asns* expression (Figure 7C). Knockdown of Atf4 also led to a general dysregulation of intracellular amino acid levels, an increase in cystine (Figure 7C) and altered GSH metabolism, such as decreased *γ*-glutamylcysteine, upon TCAi (Figure S7A and B). Overall, this work identified a key role for the ISR and Atf4 in regulating the response to TCAi and countering the associated redox and amino acid stress (Figure 7D).

**Figure 7.**
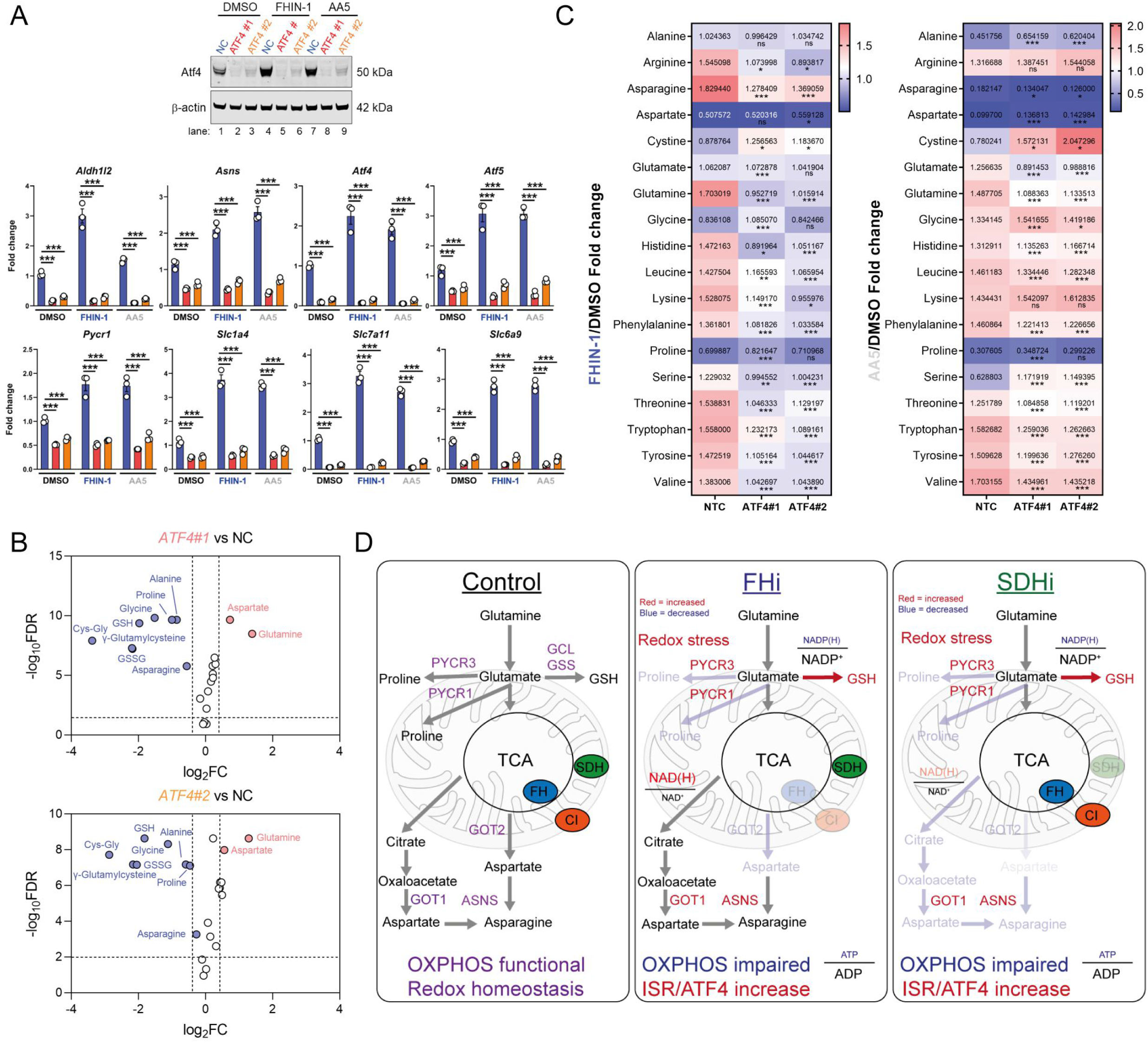
Atf4 regulates amino acid and GSH metabolism. (A) Western blot analysis of Atf4 and β-actin as loading control (24 h timepoint) (representative image of 3 independent biological replicates) and qPCR analysis of Atf4-target genes (24 h timepoint) (*n* = 3 independent biological replicates) in DMSO-, FHIN-1- and AA5-treated *Fh1^fl/fl^* cells with or without siRNA-mediated silencing of *Atf4*. Data are mean ± SEM. p value determined by ordinary one-way ANOVA, corrected for multiple comparisons using Tukey statistical hypothesis testing. p <0.05*; p <0.01**; p < 0.001***. (B) Volcano plot of glutathione-related metabolites and amino acids in Atf4-silenced *Fh1^fl/fl^* cells versus non-targeting control (NC)-*Fh1^fl/fl^* cells (48 h timepoint). (C) Heatmap of amino acid levels in FHIN-1- and AA5-versus DMSO-treated *Fh1^fl/fl^* cells with or without siRNA-mediated silencing of *Atf4* (24 h timepoint). (B-C) (*n* = 5 independent biological replicates). (D) Schematic diagram highlighting key findings of the comparative analysis between acute FHi versus SDHi.

## Discussion

Despite the importance of the TCA cycle to cellular metabolism, bioenergetics and redox homeostasis, a comprehensive and unbiased analysis of how mammalian cells respond to TCA cycle dysfunction has not been reported. The ability to perform such analysis is only now possible due to the advent of powerful mass spectrometry and next generation sequencing technologies that allow us to observe the transcriptome, metabolome and proteome in greater detail than ever before (Hasin et al., 2017). Here, we undertook a multi-modal analysis to elucidate how the TCA cycle operates and integrates with the transcriptome and proteome of the cell. From a metabolic perspective, our analysis revealed an unexpected link between a functional oxidative TCA cycle, mitochondrial respiration and thiol redox homeostasis with de novo proline biosynthesis. This is reminiscent of the work previously linking appropriate ETC function with aspartate synthesis (Birsoy *et al*., 2015; Cardaci *et al*., 2015; Sullivan *et al*., 2015) and suggests that mitochondrial function is also essential for de novo proline synthesis in certain contexts. Our analysis implicated decreased α-KG siphoning and diminished mitochondrial ATP levels as the mechanism by which the TCA cycle impairs proline synthesis. Recently, two elegant studies revealed that contrary to previous reports, NAD(P)H and not NADH is the primary redox cofactor required for mitochondrial proline synthesis (Tran *et al*., 2021; Zhu *et al*., 2021). Given the drop in the NAD(P)H/NADP^+^ ratio we observe and the reliance of mitochondrial NADP^+^ pools on ATP and the NAD kinase NADK2 (Tran *et al*., 2021; Zhu *et al*., 2021), it is tempting to speculate that a combined bioenergetic and reductive power defect plays a role upon TCAi. Furthermore, studies show that Pycr1 is a redox sensitive enzyme that can be modified by reactive electrophilic species (RES) (Timblin et al., 2021), while proline synthetic enzymes are oxidative stress-inducible (Krishnan et al., 2008) and are also targets of reactive oxygen specie (ROS)-mediated cysteine oxidation, as determined by quantitative redox proteomics of murine tissues in vivo (OxiMouse) (Xiao et al., 2020). Given TCAi leads to increased oxidative stress (and succination with FHi) and that direct perturbations of thiol redox homeostasis with mitoCDNB impair proline synthesis, it is also tempting to speculate that ROS or RES could impair this pathway. This could also explain why a common outcome of TCAi is the enhancement of GSH synthesis and why inhibiting GSH synthesis with BSO led to a further decrease in proline levels.

Intriguingly, a novel ribosome profiling technique termed diricore was recently developed to uncover signals of restrictive amino acid availability (Loayza-Puch et al., 2016). Application of diricore to a cohort of clear cell renal cell carcinoma (ccRCC) patients, a common renal cancer type associated with severe mitochondrial dysfunction and derived from kidney epithelial cells, uncovered a signal indicating proline restriction (Loayza-Puch *et al*., 2016). The authors also found that ccRCC upregulated PYCR1 in order to compensate for this proline deficit and was a targetable vulnerability in these tumours. As such, our work provides mechanistic insight into how and why proline would be restricted in the context of kidney epithelial cells with mitochondrial dysfunction and how the proline synthetic enzymes, Aldh18a1 and Pycr1, are transcriptionally regulated in this context. While proline and Pycr1 regulation has been linked to Atf4 signalling previously, this was in the context of stem cell differentiation (D’Aniello et al., 2015) and not that of mitochondrial dysfunction. Although it has been suggested that the pancreas is the major site of proline production in vivo (Tran *et al*., 2021), the kidney has been proposed to operate as a proline-producing organ under conditions of fasting and with low proline diets (Watanabe et al., 1999; Watanabe et al., 1997). Given the high density of mitochondria in proximal kidney tubules, it would be tempting to speculate that high rates of oxidative TCA activity could be important to facilitate proline biosynthesis for use by other organs under conditions of fasting or nutrient scarcity.

The importance of mitochondrial retrograde signalling to the cytosol and nucleus, and its regulation of both homeostatic and pathogenic signalling events is only beginning to be appreciated. In this study, we also investigated mitochondrial retrograde signalling events that occur in response to acute TCA cycle dysfunction to better understand the molecular underpinnings of the response. We found that TCAi increased Atf4 levels and triggered a transcriptional response reminiscent of amino deprivation and heme-deficiency. HRI has recently been implicated as the ISR kinase responsible for sensing mitochondrial dysfunction via a newly discovered pathway that involves the proteolytic cleavage of DELE1 by the protease OMA1 (Fessler et al., 2020; Guo *et al*., 2020). In contrast, asparagine has also recently been implicated in signalling mitochondrial respiration to Atf4 via GCN2, which detects uncharged aminoacyl tRNAs (Krall et al., 2021; Mick *et al*., 2020). Our data likely supports the idea that HRI is the kinase responsible for sensing TCA cycle dysfunction, however, future investigations are required to fully validate this hypothesis. Our work does however support the idea that reprogramming events are elicited to support the maintenance of both aspartate and asparagine pools under conditions of mitochondrial dysfunction. In contrast to previous studies, our analysis revealed a surprising degree of metabolic plasticity to this defect in kidney epithelial cells depending on the insult. Cells with acute FHi, but not SDHi, maintained reductive carboxylation and cytosolic aspartate synthesis likely due to decreased Complex I activity and increased NADH/NAD^+^ ratio as previously described (Birsoy *et al*., 2015; Mullen et al., 2014). However, acute FHi also led to increased consumption of the branched chain amino acids (BCAAs) valine and leucine (Figure 1I) and an increase in the cytosolic branched chain aminotransferase (BCAT1) (Figure 5D), but this was not observed with SDHi. BCAAs can act as an important nitrogen source for the synthesis of cytosolic aspartate (Mayers et al., 2016) and so another possibility for the increased capacity for cytosolic aspartate synthesis with FHi is the reprogramming of BCAA catabolism via BCAT1. Importantly, this work also suggests that the relative pool size of aspartate exceeds that of asparagine, as even under conditions of limited aspartate availability, metabolic and transcriptional changes can maintain asparagine levels under certain circumstances.

In conclusion, both FHi and SDHi impair de novo proline and aspartate synthesis by interrupting oxidative TCA cycle activity and OXPHOS (figure 7D). TCAi also decreases the NAD(P)H/NADP^+^ ratio to support cystine reduction and promote cytosolic GSH synthesis in order to counteract amino acid and redox stress. Intriguingly, acute FHi maintains asparagine synthesis via reductive carboxylation, cytosolic aspartate synthesis and an Atf4-induced increase in ASNS. However, SDHi leads to a significant impairment in asparagine synthesis as it fails to engage reductive carboxylation and promote sufficient levels of cytosolic aspartate synthesis. Finally, TCAi triggers activation of the ISR and Atf4 stabilization to communicate an amino acid-deprived and redox-stressed state to the nucleus. Overall, our work highlights the crucial role that the TCA cycle plays in maintaining metabolic homeostasis and reveals the magnificent plasticity and unique adaptations that may arise in response to metabolic perturbations.

### Limitation of the study

Although comprehensive in its metabolic investigation of acute TCA cycle dysfunction, this study was performed in a proliferating epithelial cell type of kidney origin *in vitro*. As such, whether these findings extend to different cell types or tissue types will need to be investigated further. While beyond the scope of this study, the precise mechanism(s) by which TCA cycle dysfunction triggers the ISR or how different tissues respond to TCA cycle inhibition *in vivo* under fed and fasted conditions will also be important to assess in future investigations. Finally, our study used high nutrient DMEM supplemented with 10% heat inactivated serum. The use of this medium enables careful control of our in vitro experiments and is commonly used for most molecular biology studies. Recently, it has also proved important in revealing specific metabolic dependencies relating to proline (Tran et al., 2021; Zhu *et al*., 2021). However, it will be important to assess this response under nutrient scarcity or with different nutrient compositions in the future to reveal other facets of the metabolic response.

## Methods

### Reagents and antibodies

Dimethyl sulfoxide (DMSO) D8418 (Sigma), Fumarate hydratase inhibitor 1 (FHIN-1) Cat. No.: HY-100004 (MedChemExpress), Atpenin A5 (AA5) CAS Number: 119509-24-9 (Abcam), Thenoyltrifluoroacetone (TTFA) T27006 (Sigma), mitoCDNB (Gift from Prof. Michael Murphy and Prof. Richard Hartley), U-^13^C-Glutamine (Cambridge isotopes), L-Buthionine-sulfoximine (BSO) B2515 (Sigma), ethylGSH (eGSH) G1404 (Sigma), Oligomycin A 75351 (Sigma), Anti-Puromycin antibody clone 12D10 MABE343 (Sigma), Puromycin dihydrochloride (P9620) (Sigma), Anti-Atf4 (D4B8) Rabbit mAb #11815 (CST), anti-p-eIF2α (Ser51) (D9G8) XP^®^ Rabbit mAb #3398 (CST), anti-β-Actin (13E5) Rabbit mAb #4970 (CST), Integrated stress response inhibitor (ISRIB) SML0843 (Sigma), ^35^S-methionine NEG009L005MC (PerkinElmer), Glucose (Sigma), Glutamine (Sigma), Pyruvate (Sigma), Carbonyl cyanide-p-trifluoromethoxyphenylhydrazone (FCCP) C2920 (Sigma), Antimycin A A8674 (Sigma), Rotenone R8875 (Sigma), Silencer^®^ select Non-targeting control (NTC) siRNA Catalog # 4390843 (ThermoFisher Scientific), Silencer^®^ select *Atf4* siRNA #1 Catalog # 4390771 Assay ID s62691(Thermo Scientific), Silencer^®^ select^™^ *Atf4* siRNA #2 Catalog # 4390771 Assay ID s62690 (Thermo Scientific), Lipofectamine^®^ RNAiMAX Catalog # 13778075 (ThermoFisher Scientific)

### Drug treatments

All compounds used DMSO as a vehicle except eGSH, which used ultrapure water. All compounds were made up as stock concentrations and diluted 1/1000 – 1/500 in cell culture medium (CCM) prior to treatment. CCM with treatment compounds were vortexed briefly to ensure even distribution of the compounds before adding to appropriate wells or dishes for the indicated timepoints (1, 3, 6 or 24 h). The final percentage of DMSO in CCM was never more than 0.2%.

### Cell culture

*Fh1*-proficient (*Fh1^fl/fl^*) cells and the two *Fh1*-deficient clones (*Fh1*^-/-*CL1*^ and *Fh1*^-/-*CL19*^) were obtained, as previously described (Frezza et al., 2011; Sciacovelli et al., 2016). *Fh1*^-/-*pFlh-GFP*^ cells were generated from *Fh1*^-/-*CL1*^ after stable expression of a plasmid carrying either fulllength, as previously described (Sciacovelli *et al*., 2016). *Sdhb*-proficient (*Sdhb^fl/fl^*) and two *Sdhb*-deficient clones (*Sdhb*^-/-*CL5*^ and *Sdhb*^-/-*CL7*^) were a gift from Prof. Eyal Gottlieb (Cardaci *et al*., 2015). Murine cells were cultured using DMEM (Gibco-41966-029) supplemented with 10% heat-inactivated serum (Gibco-10270-106). Genotyping of cells was assessed as previously described (Sciacovelli *et al*., 2016). All cells were authenticated by short tandem repeat (STR) and routinely checked for mycoplasma contamination. Counting for cell plating and volume measurement were obtained using the CASY cell counter (Omni Life Sciences). Briefly, cells were gently detached using trypsin-EDTA 0.05%, centrifuged at 1500 rpm for 5 mins and the cell pellet was re-suspended in fresh media prior to counting.

### LC-MS Metabolomics

#### Steady-state metabolomics

For steady-state metabolomics, 5×10^4^ cells were plated the day before onto 6-well plates (5 biological replicates from independently maintained cell lines) and extracted at the appropriate experimental endpoint. Prior to metabolite extraction, cells were counted using CASY cell counter (Omni Life Sciences) using a separate counting plate prepared in parallel and treated exactly like the experimental plate. For consumption release experiments (CoRe) experiments a Day 0 counting plate was also required to assess proliferation rates. At the experimental endpoint, an aliquot of cell culture conditioned media (CCM) was collected to facilitate CoRe analysis. The remaining media was aspirated off and the cells were washed at room temperature with PBS and placed on a cold bath with dry ice. Metabolite extraction buffer (MES) was added to each well following the proportion 1×10^6^ cells/0.5 ml of buffer. After 10 minutes, the plates were stored at −80°C freezer and kept overnight. The following day, the extracts were scraped and mixed at 4°C for 15 min in a thermomixer at 2000 rpm. After final centrifugation at max speed for 20 min at 4°C, the supernatants were transferred into labelled LC-MS vials.

#### Tracing experiments

5×10^4^ cells were plated onto 6-well plate (5 biological replicates from independently maintained cell lines for each condition). At the experimental starting point, the medium was replaced with fresh media containing fully labelled U-^13^C-Glutamine (obtained from Cambridge Isotopes Laboratories) and the compound(s) of interest and left on for the duration of the experiment (24 h)

#### Liquid chromatography coupled to Mass Spectrometry (LC-MS) analysis

HILIC chromatographic separation of metabolites was achieved using a Millipore Sequant ZIC-pHILIC analytical column (5 μm, 2.1 × 150 mm) equipped with a 2.1 × 20 mm guard column (both 5 mm particle size) with a binary solvent system. Solvent A was 20 mM ammonium carbonate, 0.05% ammonium hydroxide; Solvent B was acetonitrile. The column oven and autosampler tray were held at 40 °C and 4 °C, respectively. The chromatographic gradient was run at a flow rate of 0.200 mL/min as follows: 0–2 min: 80% B; 2-17 min: linear gradient from 80% B to 20% B; 17-17.1 min: linear gradient from 20% B to 80% B; 17.1-22.5 min: hold at 80% B. Samples were randomized and analyzed with LC–MS in a blinded manner with an injection volume was 5 μl. Pooled samples were generated from an equal mixture of all individual samples and analyzed interspersed at regular intervals within sample sequence as a quality control.

Metabolites were measured with a Thermo Scientific Q Exactive Hybrid Quadrupole-Orbitrap Mass spectrometer (HRMS) coupled to a Dionex Ultimate 3000 UHPLC. The mass spectrometer was operated in full-scan, polarity-switching mode, with the spray voltage set to +4.5 kV/-3.5 kV, the heated capillary held at 320 °C, and the auxiliary gas heater held at 280 °C. The sheath gas flow was set to 25 units, the auxiliary gas flow was set to 15 units, and the sweep gas flow was set to 0 unit. HRMS data acquisition was performed in a range of *m/z* = 70–900, with the resolution set at 70,000, the AGC target at 1 × 10^6^, and the maximum injection time (Max IT) at 120 ms. Metabolite identities were confirmed using two parameters: (1) precursor ion m/z was matched within 5 ppm of theoretical mass predicted by the chemical formula; (2) the retention time of metabolites was within 5% of the retention time of a purified standard run with the same chromatographic method. Chromatogram review and peak area integration were performed using the Thermo Fisher software Tracefinder 5.0 and the peak area for each detected metabolite was normalized against the total ion count (TIC) of that sample to correct any variations introduced from sample handling through instrument analysis. The normalized areas were used as variables for further statistical data analysis.

For ^13^C-tracing analysis, the theoretical masses of ^13^C isotopes were calculated and added to a library of predicted isotopes. These masses were then searched with a 5 ppm tolerance and integrated only if the peak apex showed less than 1% difference in retention time from the [U-^12^C] monoisotopic mass in the same chromatogram. After analysis of the raw data, natural isotope abundances were corrected using the AccuCor algorithm (https://github.com/lparsons/accucor).

### RNA sequencing

5×10^4^ cells were plated onto 3 replicate 6-cm dishes before the extraction and treated as indicated. RNA isolation was carried using RNeasy kit (Qiagen) following manufacturer’s suggestions and eluted RNA was purified using RNA Clean & Concentrator Kits (Zymo Research). RNA-seq samples libraries were prepared using TruSeq Stranded mRNA (Illumina) following the manufacturer’s description. For the sequencing, the NextSeq 75 cycle high output kit (Illumina) was used and samples spiked in with 1% PhiX. The samples were run using NextSeq 500 sequencer (Illumina). Differential Gene Expression Analysis was done using the counted reads and the R package edgeR version 3.26.5 (R version 3.6.1) for the pairwise comparisons.

### Cellular fractionation and mitochondrial isolation

Cells were grown to confluency on 2 × 150 mm dishes per indicated treatments. At the experimental endpoint, cells were washed 4 × with ice cold 1X PBS on ice. All excess PBS was removed after the final wash. 500 μl of ice-cold STE buffer (250 mM sucrose, 5 mM Tris, 1 mM EGTA, pH 7.4, 4°C) was added to each dish and cells were scraped and transferred to a pre-cooled 7 ml dounce homogenizer and cells homogenized with ~100 strokes of a tight-fitting pestle. The homogenate was passed through a 30G needle ten times and split into 1.5 mL centrifuge tubes. Cells were then centrifuged at 3,000 × g for 3 mins at 4°C. The supernatant was transferred to a new centrifuge tube and centrifuged as before. The supernatant was transferred to a new tube and centrifuge at 11,000 × g for 5 mins at 4°C. The supernatant ‘cytosol’ fraction was taken and placed in a new centrifuge tube prior to metabolomic extraction as above. The remaining pellet was resuspended in 1 mL STE buffer and centrifuged again for 11,000 × g for 5 mins at 4°C to generate the ‘mitochondrial’ fraction. All supernatant was removed prior to metabolite extraction.

### Proteomic analysis

#### Sample preparation

5×10^4^ cells were plated onto 5 replicate 10-cm dishes, grown to confluence and treated as indicated. At the experimental endpoint, cells were washed with PBS on ice and centrifuged at 1500 rpm for 5 mins at 4°C and frozen at −80°C. Cell pellets were lysed, reduced and alkylated in 100 μl of 6M Gu-HCl, 200 mM Tris-HCl pH 8.5, 1 mM TCEP, 1.5 mM Chloroacetamide by probe sonication and heating to 95°C for 5 min. Protein concentration was measured by a Bradford assay and initially digested with LysC (Wako) with an enzyme to substrate ratio of 1/200 for 4 h at 37 °C. Subsequently, the samples were diluted 10fold with water and digested with porcine trypsin (Promega) at 37°C overnight. Samples were acidified to 1% TFA, cleared by centrifugation (16,000 g at RT) and approximately 20 μg of the sample was desalted using a Stage-tip. Eluted peptides were lyophilized, resuspended in 0.1% TFA/water and the peptide concentration was measured by A280 on a nanodrop instrument (Thermo). The sample was diluted to 1 μg/ 5 μl for subsequent analysis.

#### Mass spectrometry analysis

The tryptic peptides were analyzed on a Fusion Lumos mass spectrometer connected to an Ultimate Ultra3000 chromatography system (both Thermo Scientific, Germany) incorporating an autosampler. 5 μL of the tryptic peptides, for each sample, was loaded on an Aurora column (Ionoptiks, Melbourne Australia) and separated by an increasing acetonitrile gradient, using a 150-min reverse-phase gradient (from 3%–40% Acetonitrile) at a flow rate of 400 nL/min. The mass spectrometer was operated in positive ion mode with a capillary temperature of 220°C, with a potential of 1500 V applied to the column. Data were acquired with the mass spectrometer operating in automatic data-dependent switching mode, with MS resolution of 240k, with a cycle time of 1 s and MS/MS HCD fragmentation/analysis performed in the ion trap. Mass spectra were analyzed using the MaxQuant Software package in biological triplicate. Label-free quantitation was performed using MaxQuant. All the samples were analyzed as biological triplicates.

#### Data analysis

Data were analyzed using the MaxQuant software package. Raw data files were searched against a human database (Uniprot Homo sapiens), using a mass accuracy of 42.5 ppm and 0.01 false discovery rate (FDR) at both peptide and protein level. Every single file was considered as separate in the experimental design; the replicates of each condition were grouped for the subsequent statistical analysis. Carbamidomethylation was specified as fixed modification while methionine oxidation and acetylation of protein N-termini were specified as variable. Subsequently, missing values were replaced by a normal distribution (1.8 π shifted with a distribution of 0.3 π) in order to allow the following statistical analysis. Results were cleaned for reverse and contaminants and a list of significant changes was determined based on average ratio and t-test. LFQ-analyst was used to perform differential expression analysis after pre-processing with MaxQuant (Shah et al., 2020).

### Oxygen consumption rate (OCR) measurements

Cellular respiration (OCR), as part of the Cell Mito Stress test, was measured using the real-time flux analyzer XF-24e Seahorse (Agilent). In brief, 6×10^4^ cells were plated onto the instrument cell plate 24 h before the experiment in complete DMEM with 10 % FBS (5 replicate wells for each condition). The following day cells were treated as indicated for 24 h. At the treatment endpoint, the medium was replaced with DMEM pH 7.4 (Agilent) supplemented with 25 mM glucose, 2.5 mM glutamine and 1 mM pyruvate prior to Cell Mito Stress Test analysis according to manufacturer’s instructions. Cells were treated with 2.5 μM Oligomycin, 0.5μM FCCP and 0.5 μM Antimycin A/Rotenone to assess different respiration parameters.

### RNA extraction and real-time quantitative (qPCR)

5×10^4^ cells were plated onto 6-well plates, left to adhere overnight and treated for indicated times. At experimental endpoint, cells were washed in 1X PBS and then RNA was extracted using RNeasy kit (Qiagen) following the manufacturer’s instructions. RNA was eluted in water and then quantified using a Nanodrop (ThermoFisher Scientific). 1 μg of RNA was reverse-transcribed using High capacity cDNA Reverse Transcription kit (Applied Biosystems). For real-time qPCR, cDNA was run using Fast SYBR^®^ Green Master Mix (Applied Biosystems) according to manufacturer’s instructions and primers were designed for genes of interest (see below) using primer-BLAST. *Actb* was used as the endogenous control. qPCR experiments were run on a 384-well QuantStudio^™^ Real-Time PCR system (ThermoFisher Scientific).

### qPCR primers

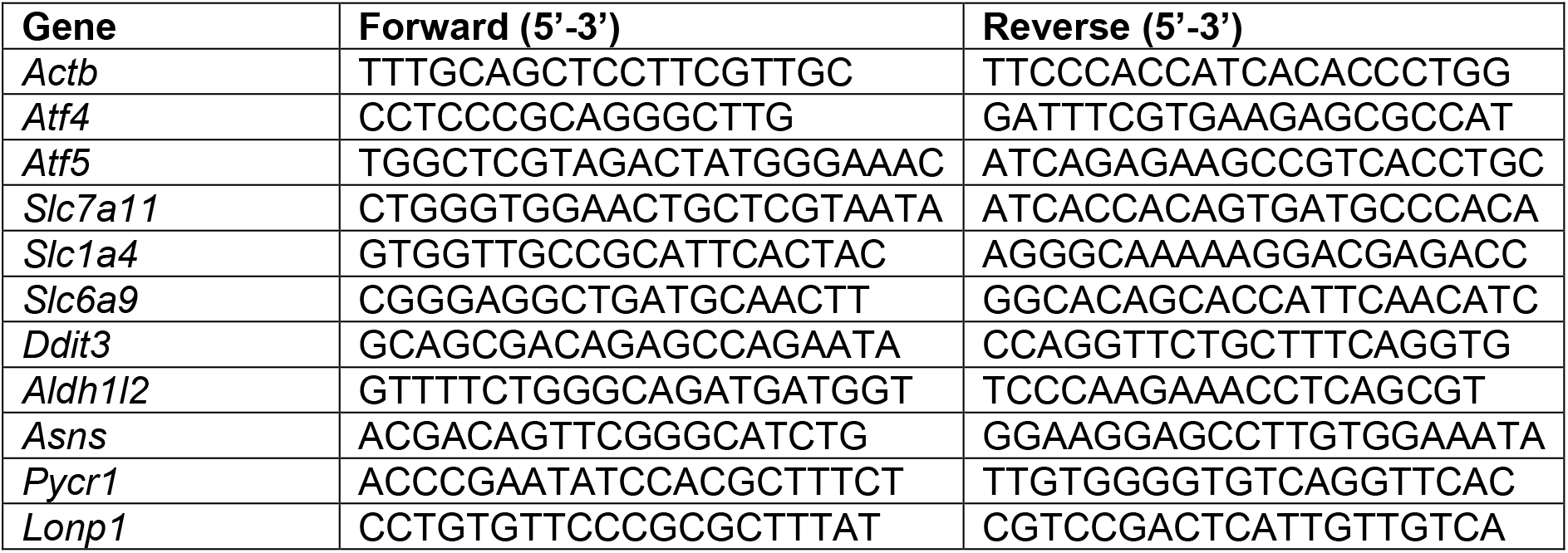

### Puromycin assay

5×10^4^ cells were plated onto 12-well plates, left to adhere overnight and treated with FHIN-1, AA5 and TTFA for indicated times (1, 3 or 6 h). For the last 15 minutes of the treatment time, 20μg/mL puromycin dihydrochloride (Sigma) was added to the well prior to extraction. Puromycin incorporation into neosynthesized proteins was assessd with Western blotting and an anti-puromycin antibody (Sigma).

### Western blotting

5×10^4^ cells were plated onto 6- or 12-well plates, left to adhere overnight and treated for indicated times. At the experimental endpoint, cells were counted on a parallel counting plate using a CASY counter, washed in 1X PBS and then lysed on ice with the appropriate volume (100 μL per 100,000 cells) of 4X Bolt^™^ Loading buffer (Thermo Scientific) diluted to 1X with RIPA buffer (150mM NaCl, 1%NP-40, Sodium deoxycholate (DOC) 0.5%, sodium dodecyl phosphate (SDS) 0.1%, 25mM Tris) supplemented with protease and phosphates inhibitors (Protease inhibitor cocktail, Phosphatase inhibitor cocktail 2/3) (Sigma-Aldrich) and containing 4% β-mercaptoethanol. Protein samples were then heated at 70°C for 10 minutes, briefly centrifuged and stored at −20C for future use. Samples were loaded onto 4-12% Bis-Tris Bolt^™^ gradient gels and run at 160V constant for 1h in Bolt^™^ MES 1X running buffer (Thermo Scientific). Dry transfer of the proteins onto a nitrocellulose membrane was done using iBLOT2 (Thermo Scientific) for 12 minutes at 20V. Membranes were incubated in blocking buffer for 1h (either 5% BSA or 5% milk in TBS 1X +0.01 % Tween-20, TBST 1X). Primary antibodies were incubated in blocking buffer ON at 4°C under gentle agitation. On the following day, the membranes were washed three times in TBST 1X for 5 mins and then secondary antibodies (conjugated with 680 or 800 nm fluorophores) (Li-Cor) incubated for 1 h at room temperature at 1:2000 dilution in blocking buffer. Images were acquired and quantified using Image Studio lite 5.2 (Li-Cor) on Odyssey CLx instrument (Li-Cor).

### RNAi transfection

5×10^4^ cells were plated onto 6-well plates and left to adhere overnight. On the following day, plates were transfected with indicated Silencer^®^ select siRNAs (10 nM) complexed with Lipofectamine^®^ RNAiMAX and Opti-MEM^®^ medium according to manufacturer’s instructions. After 24 h incubation, cells were treated as indicated prior to downstream processing for metabolomics, Western blot or qPCR.

### ^35^S-methionine labelling of mitochondrial translation products

In order to label newly synthesised mitochondrially expressed proteins, the previously published protocol was used (Pearce et al., 2017). Briefly, cells at approximately 80% confluency were incubated in methionine/cysteine-free medium for 10 min before the medium was replaced with methionine/cysteine-free medium containing 10% dialysed FCS and emetine dihydrochloride (100 μg/ml) to inhibit cytosolic translation. Following a 20 min incubation, 120 μCi/ml of [^35^S]-methionine was added and the cells were incubated for 30 min. After washing with 1X PBS, cells were lysed, and 30 μg of protein was loaded on 10–20% Tris-glycine SDS-PAGE gels. Dried gels were visualized with a PhosphorImager system.

### Statistical analysis

Graphs were generated using Graphpad prism 9.0 software, which was used to perform most statistical analysis. Metaboanalyst 5.0 was used to analyze metabolomics data from Figure 1 A-C. ORA analysis of the significant hits from RNAseq and proteomics used Enrichr (Chen *et al*., 2013). GSEA analysis of RNAseq was performed using the Broad Institutes GSEA 4.1.0 (Subramanian *et al*., 2005).

## Author contributions

D.G.R and C.F conceptualized the study. D.G.R designed and performed most of the experiments and analysis, interpreted the data and coordinated the research. D.G.R prepared the figures and wrote the manuscript with assistance from all other authors. M.Y. and E.N. ran and analyzed the metabolomics samples. H.A.P. performed fractionation experiments and provided important advice and intellectual input. G.R.B and A.v.K performed proteomic sample preparation and analysis. C.A.P performed 35S-methionine labelling of mitochondrial translation products under the supervision of M.M. M.S performed Antimycin A experiment. T.Y, N.B and J.L.M aided with data generation. M.P.M provided reagents and advice. C.F. obtained funding, edited the manuscript and oversaw the research programme.

## Acknowledgements

We thank Cambridge Genomic Services (Department of Pathology, University of Cambridge) especially Dr Alexandria Karcanias and Dr Julien Bauer for the RNA-seq library preparation, sequencing and differential expression analysis. We also thank all the Frezza lab members for all their helpful discussions and insights and Prof. Richard Hartley who originally synthesized mitoCDNB. M.M and C.A.P are funded by Medical Research Council (MRC) core grant to the MRC Mitochondrial Biology Unit (MC_UU_00015/4).

---

**Figure S1.**
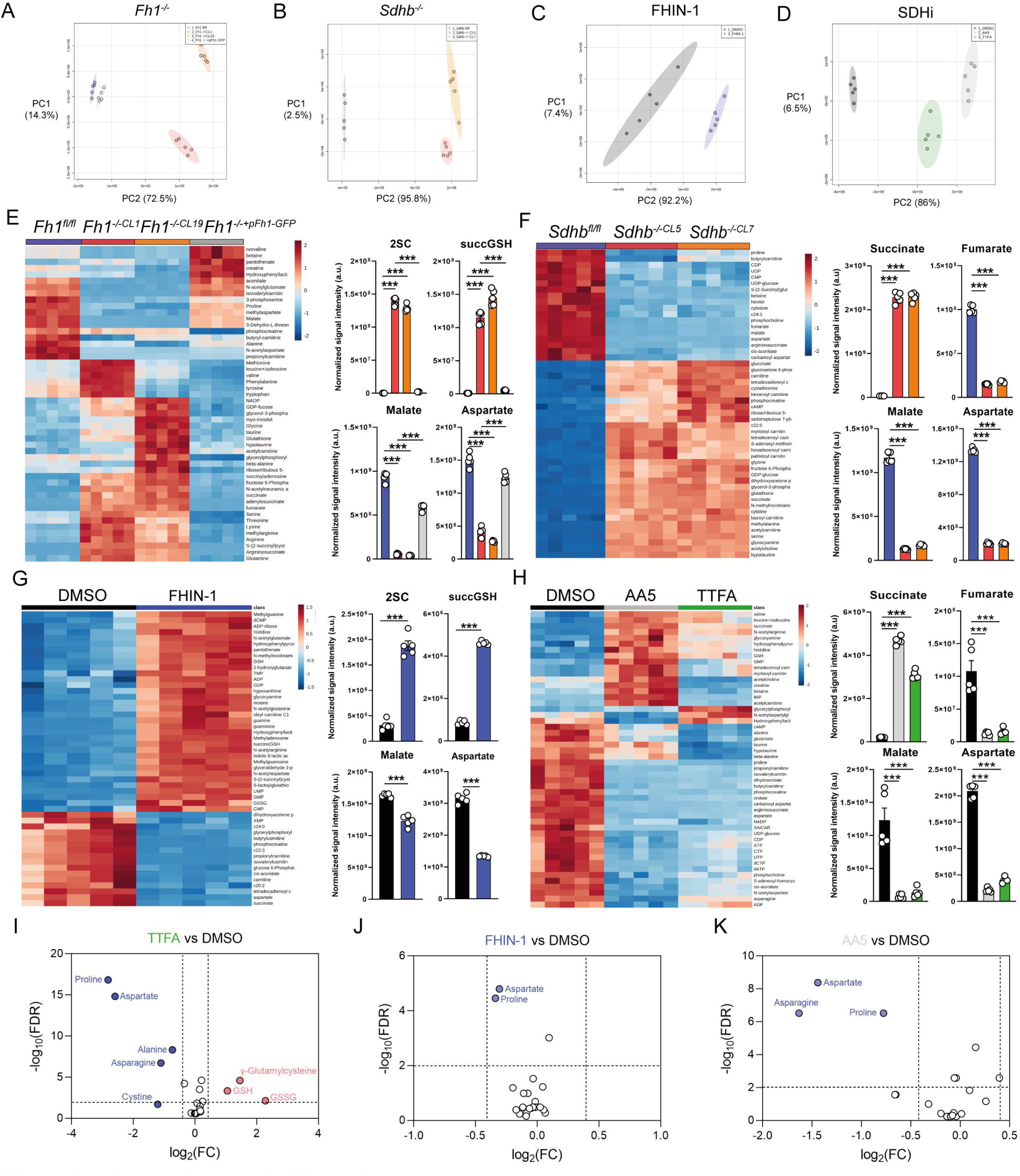
Supporting data for figure 1. (A) PCA plot of metabolomics data of *Fh1^fl/fl^*, *Fh1*^-/-*CL1*^, *Fh1*^-/-*CL19*^ and *Fh1*^-/-*CL1*+*pFh1*^ cells (*n* = 5 independent biological replicates). (B) PCA plot of metabolomics data of *Sdhb^fl/fl^*, *Sdhb*^-/-*CL5*^ and *Sdhb*^-/-*CL7*^ cells (*n* = 5 independent biological replicates). (C) PCA plot of DMSO- and FHIN-1 (20 μM)-treated *Fh1^fl/fl^* cells (24 h timepoint) (*n* = 5 independent biological replicates). (D) PCA plot of DMSO-, AA5 (1 μM)- and TTFA (500 μM)-treated *Fh1^fl/fl^* cells (24 h timepoint) (*n* = 5 independent biological replicates). (E) Heatmap of top 50 significant metabolite changes in *Fh1^fl/fl^*, *Fh1*^-/-*CL1*^, *Fh1*^-/-*CL19*^ and *Fh1*^-/-*CL1*+*pFh1*^ cells (*n* = 5 independent biological replicates) and relative change in *Fh1*-deficient metabolic markers (S)-2-succinocysteine (2SC), succinicGSH, malate and aspartate (*n* = 5 independent biological replicates). (F) Heatmap of top 50 significant metabolite changes in *Sdhb^fl/fl^*, *Sdhb*^-/-*CL5*^ and *Sdhb*^-/-*CL7*^ cells (*n* = 5 independent biological replicates) and relative change in *Sdh*-deficient metabolic markers succinate, fumarate, malate and aspartate (*n* = 5 independent biological replicates). (G) Heatmap of top 50 significant metabolite changes in DMSO- and FHIN-1 (20 μM)-treated *Fh1^fl/fl^* cells (24 h timepoint) (*n* = 5 independent biological replicates) and relative change in *Fh1*-deficient metabolic markers 2SC, succinicGSH, malate and aspartate (*n* = 5 independent biological replicates). (H) Heatmap of top 50 significant metabolite changes in DMSO-, AA5 (1 μM)- and TTFA (500 μM)-treated *Fh1^fl/fl^* cells (24 h timepoint) (*n* = 5 independent biological replicates) and relative change in *Sdh*-deficient metabolic markers succinate, fumarate, malate and aspartate (*n* = 5 independent biological replicates). (E-H) Data are mean ± SEM. p value determined by ordinary one-way ANOVA, corrected for multiple comparisons using Tukey statistical hypothesis testing. p <0.05*; p <0.01**; p < 0.001***. (I) Volcano plot of log2 fold change in glutathione-related metabolites and amino acids in TTFA (500 μM)-treated *Fh1^fl/fl^* versus DMSO-vehicle control *Fh1*^fl/fl^ cells (24 h timepoint) (*n* = 10 independent biological replicates). (J) Volcano plot of log2 fold change in glutathione-related metabolites and amino acids in FHIN-1 (20 μM)-treated *Fh1^fl/fl^* versus DMSO-vehicle control *Fh1^fl/fl^* cells (1 h timepoint) (*n* = 5 independent biological replicates). (K) Volcano plot of log2 fold change glutathione-related metabolites and amino acids in AA5 (1 μM)-treated *Fh1*^fl/fl^ versus DMSO-treated *Fh1*^fl/fl^ cells (1 h timepoint) (*n* = 5 independent biological replicates). p value determined by multiple unpaired t-tests, corrected with two-stage step-up method of Benjamini, Krieger and Yekutieli - FDR = 5%.

**Figure S2.**
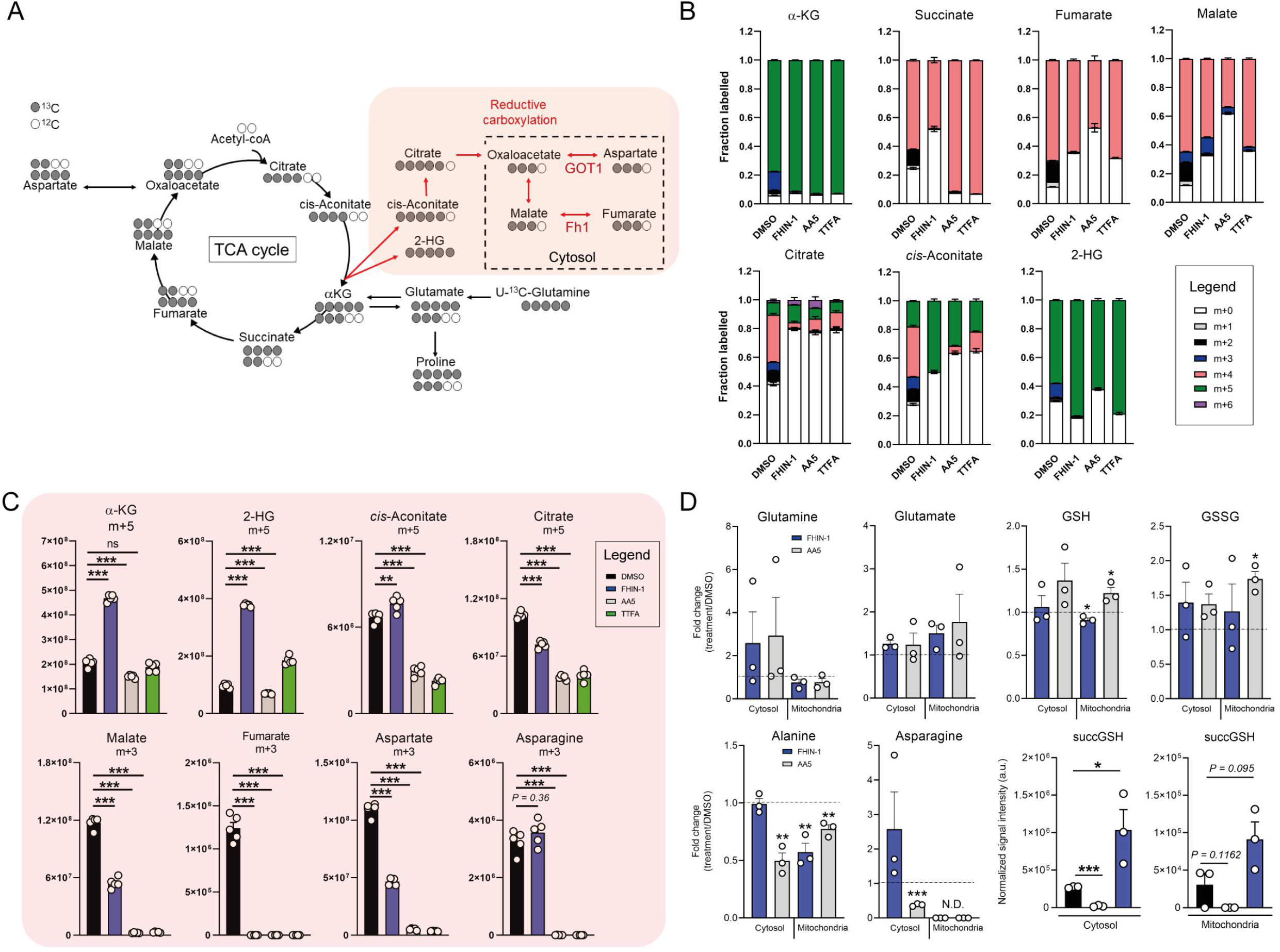
Supporting data for figure 2. (A) Schematic diagram highlighting U-^13^C-glutamine tracing into the TCA cycle and reductive carboxylation. (B) U-^13^C-glutamine tracing into α-KG, succinate, fumarate, malate, citrate, cis-aconitate and 2-HG (total isotopologue fraction distribution) in DMSO, FHIN-1 (20 μM), AA5 (1 μM) and TTFA (500 μM)-treated *Fh1^fl/fl^* cells (24 h timepoint) (*n* = 5 independent biological replicates). (C) U-^13^C-glutamine tracing into α-KG, succinate, fumarate, malate, citrate, cis-aconitate and 2-HG (m+5 and m+3 labelling intensity) in DMSO, FHIN-1 (20 μM), AA5 (1 μM) and TTFA (500 μM)-treated *Fh1^fl/fl^* cells (24 h timepoint) (*n* = 5 independent biological replicates). Data are mean ± SEM. p value determined by ordinary one-way ANOVA, corrected for multiple comparisons using Tukey statistical hypothesis testing. p <0.05*; p <0.01**; p < 0.001***. (D) Glutamine, glutamate, GSH, GSSG, alanine, asparagine level fold changes and succGSH relative intensity in mitochondrial and cytosol fractions in FHIN-1 (20 μM)- and AA5(1 μM)-treated versus DMSO-treated *Fh1^fl/fl^* cells (24 h timepoint) (*n* = 3 independent biological replicates). Data are mean ± SEM. p values determined by unpaired two-tailed t-test. p <0.05*; p <0.01**; p < 0.001***.

**Figure S3.**
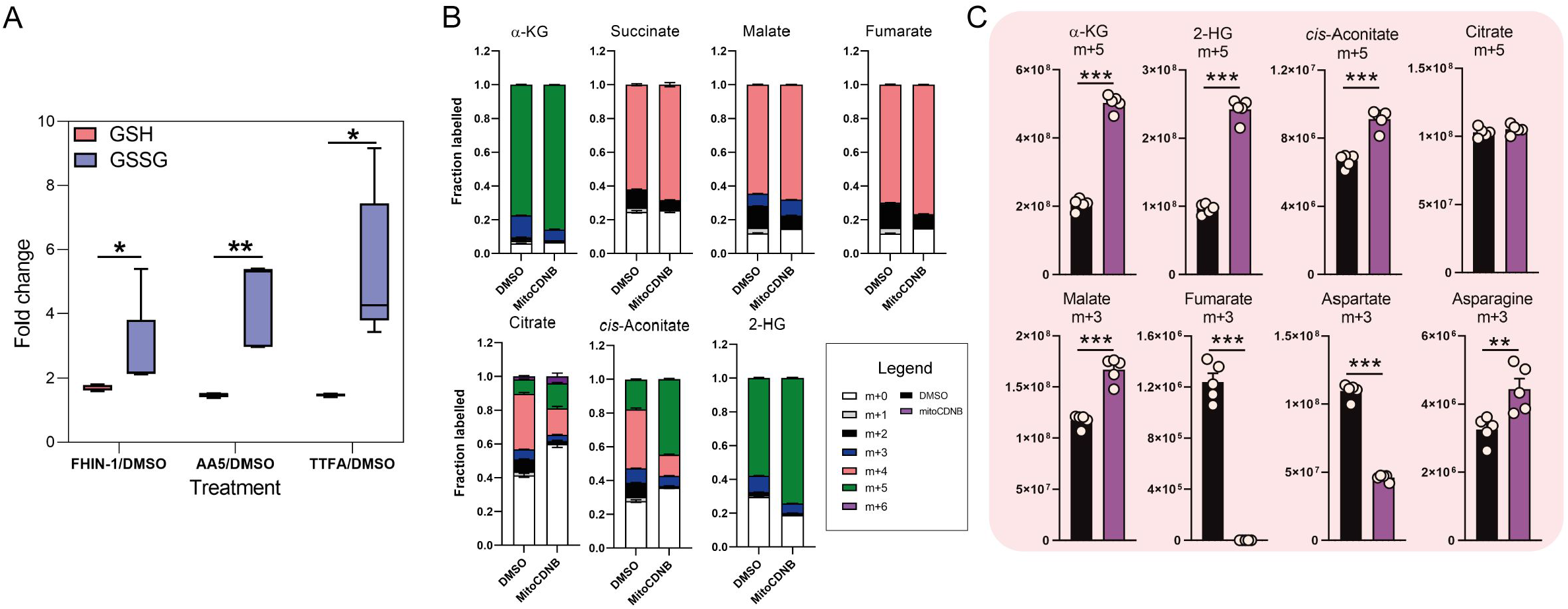
Supporting data for figure 3. (A) Interleaved box and whiskers plot of fold change in GSH and GSSG with FHIN-1 (20 μM)-, AA5 (1 μM)- and TTFA (500 μM)-treated *Fh1^fl/fl^* cells versus DMSO-treated *Fh1^fl/fl^* cells (24 h timepoint) (*n* = 5 independent biological replicates). p values determined by unpaired two-tailed t-test. p <0.05*; p <0.01**; p < 0.001***. (B) U-^13^C-glutamine tracing into α-KG, succinate, fumarate, malate, citrate, cis-aconitate and 2-HG (total isotopologue fraction distribution) in DMSO and mitoCDNB (10 μM)-treated *Fh1^fl/fl^* cells (24 h timepoint) (*n* = 5 independent biological replicates). (C) U-^13^C-glutamine tracing into α-KG, succinate, fumarate, malate, citrate, cis-aconitate and 2-HG (m+5 and m+3 labelling intensity) in DMSO and mitoCDNB (10 μM)-treated *Fh1^fl/fl^* cells (24 h timepoint) (*n* = 5 independent biological replicates). Data are mean ± SEM. p values determined by unpaired two-tailed t-test. p <0.05*; p <0.01**; p < 0.001***.

**Figure S4.**
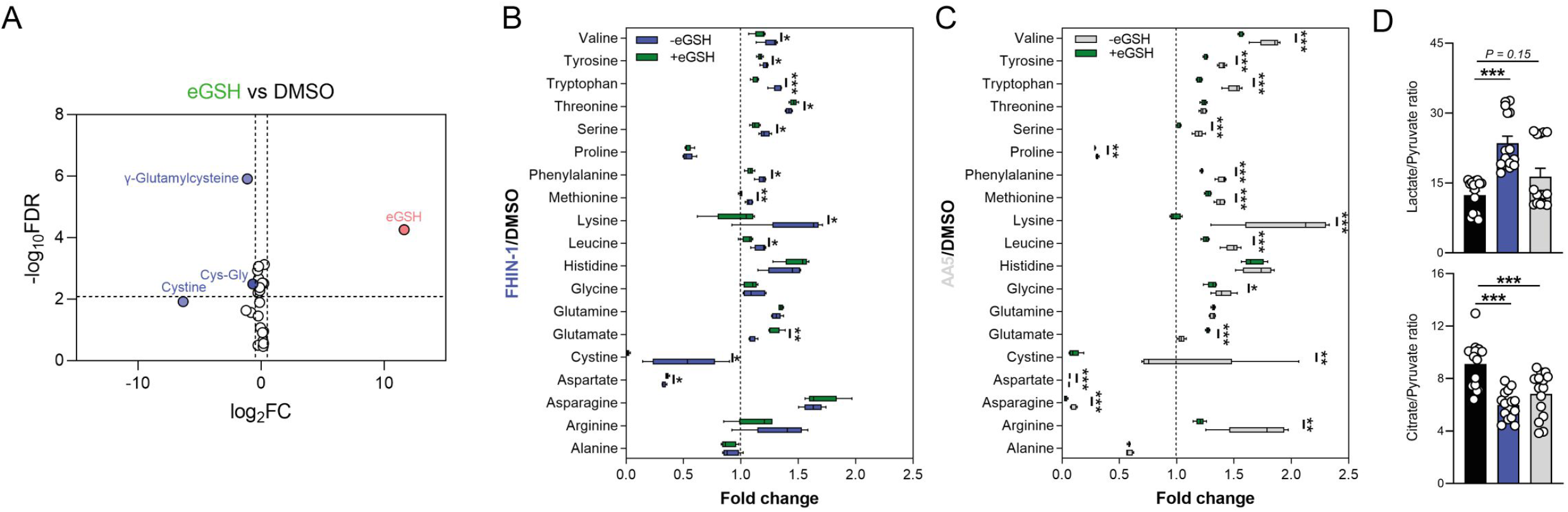
Supporting data for figure 4. (A) Volcano plot of log2 fold change in glutathione-related metabolites and amino acids in eGSH (1 mM)-treated versus DMSO-treated *Fh1*^fl/fl^ cells (24 h timepoint) (*n* = 5 independent biological replicates). (B) Interleaved box and whiskers plot of fold change in amino acids with FHIN-1 (20 μM)-treated or FHIN-1 (20 μM) and eGSH (1 mM) co-treatment of *Fh1^fl/fl^* cells (24 h timepoint) (*n* = 5 independent biological replicates). (C) Interleaved box and whiskers plot of fold change in amino acids with AA5 (1 μM)-treated or AA5 (1 μM) and eGSH (1 mM) co-treatment of *Fh1^fl/fl^* cells (24 h timepoint) (*n* = 5 independent biological replicates). (A-C) p value determined by multiple unpaired t-tests, corrected with two-stage step-up method of Benjamini, Krieger and Yekutieli - FDR = 5%. p <0.05*; p <0.01**; p < 0.001***. (D) Lactate/pyruvate and citrate/pyruvate ratio in DMSO-, FHIN-1 (20 μM)- and AA5 (1 μM)-treated *Fh1^fl/fl^* cells (24 h timepoint) (*n* = 15 independent biological replicates). Data are mean ± SEM. p values determined by unpaired two-tailed t-test. p <0.05*; p <0.01**; p < 0.001***.

**Figure S5.**
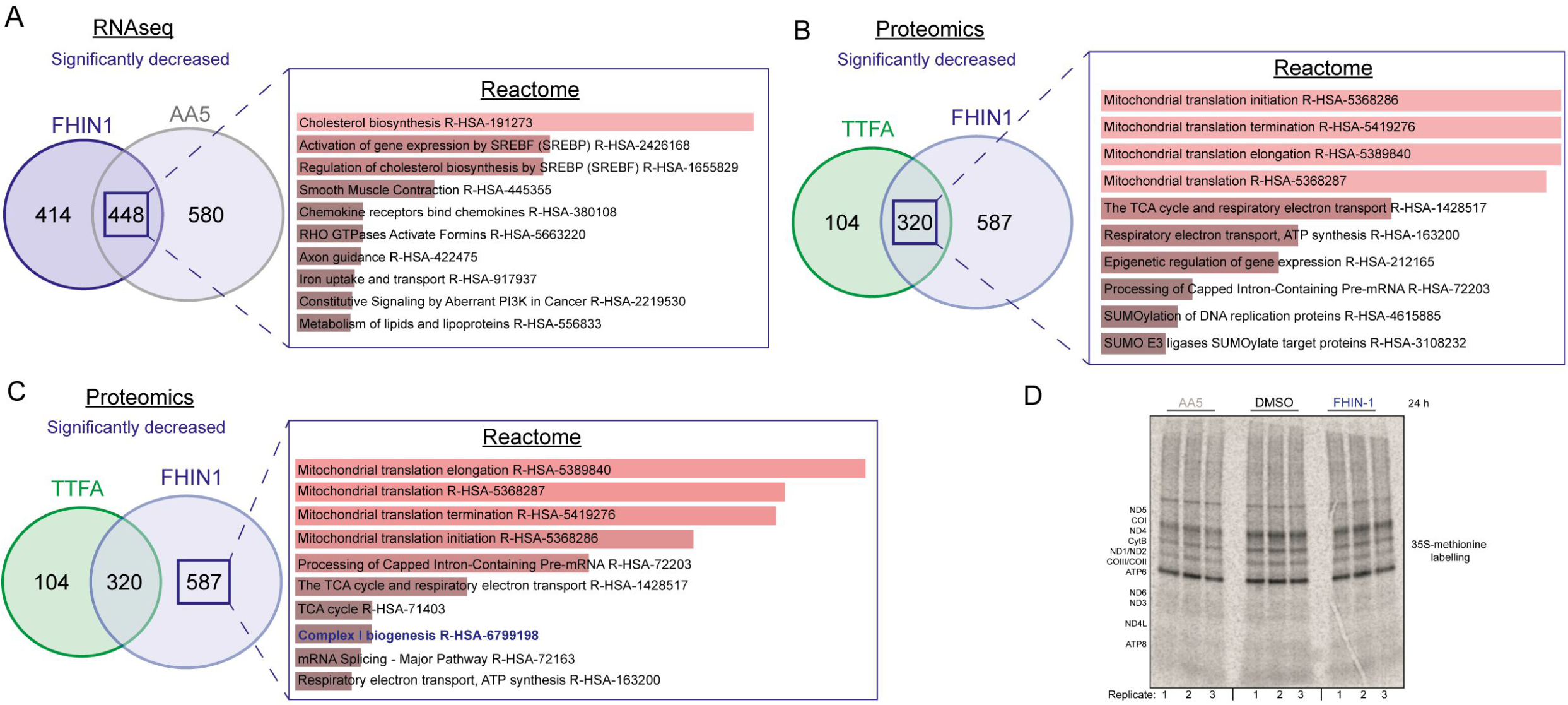
Supporting data for figure 5. (A) ORA of overlapping transcripts significantly decreased in FHIN-1 (20 μM)- and AA5 (1 μM)-treated *Fh1^fl/fl^* cells (24 h timepoint) (24 h timepoint) (*n* = 3 independent biological replicates). (B) ORA of overlapping proteins significantly decreased in FHIN-1 and TTFA-treated *Fh1^fl/fl^* cells (24 h timepoint) (*n* = 5 independent biological replicates). (C) ORA of proteins significantly decreased uniquely in FHIN-1-treated *Fh1^fl/fl^* cells (24 h timepoint) (*n* = 5 independent biological replicates).(A-C) Reactome pathways ranked by combined score (p value and Z score). (D) 35S-methionine labelling of mitochondrial translation in DMSO-, FHIN-1 (20 μM)- and AA5 (1 μM)-treated *Fh1^fl/fl^* cells (24 h timepoint) (*n* = 3 independent biological replicates).

**Figure S6.**
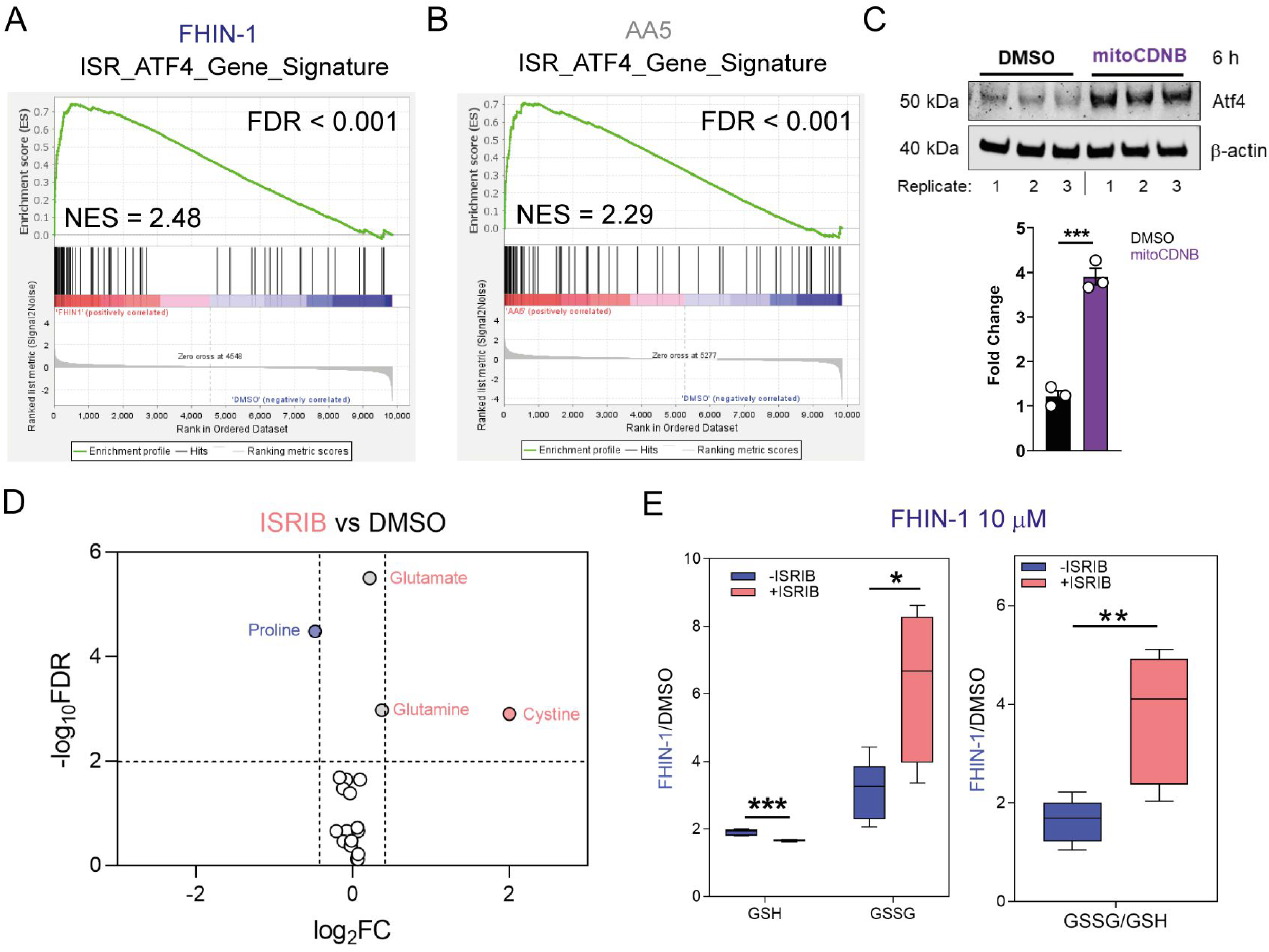
Supporting data for figure 6. (A) GSEA of Atf4 gene signature queried with molecular signature database (MsigDB) hallmarks geneset on RNA sequencing data from FHIN-1 (20 μM)-versus DMSO-treated *Fh1^fl/fl^* cells (24 h timepoint) (*n* = 3 independent biological replicates). (B) GSEA of Atf4 gene signature queried with MsigDB hallmarks geneset on RNA sequencing data from AA5 (1 μM)-versus DMSO-treated *Fh1^fl/fl^* cells (24 h timepoint) (*n* = 3 independent biological replicates). (C) Western blot analysis and quantification of Atf4 and β-actin as loading control in DMSO and mitoCDNB (10 μM)-treated *Fh1^fl/fl^* cells (6 h timepoint) (*n* = 3 independent biological replicates). Data are mean ± SEM. p values determined by unpaired two-tailed t-test. p <0.05*; p <0.01**; p < 0.001***. (D) Volcano plot of log2 fold change in glutathione-related metabolites and amino acids in ISRIB (500 nM)-treated versus DMSO-treated *Fh1*^fl/fl^ cells (24 h timepoint) (*n* = 5 independent biological replicates). (E) Interleaved box and whiskers plot of fold change in GSH and GSSG in FHIN-1 (10 μM)-treated or FHIN-1 and ISRIB co-treated *Fh1^fl/fl^* cells and the ratio of the fold change (24 h timepoint) (*n* = 5 independent biological replicates). p values determined by unpaired two-tailed t-test. p <0.05*; p <0.01**; p < 0.001***.

**Figure S7.**
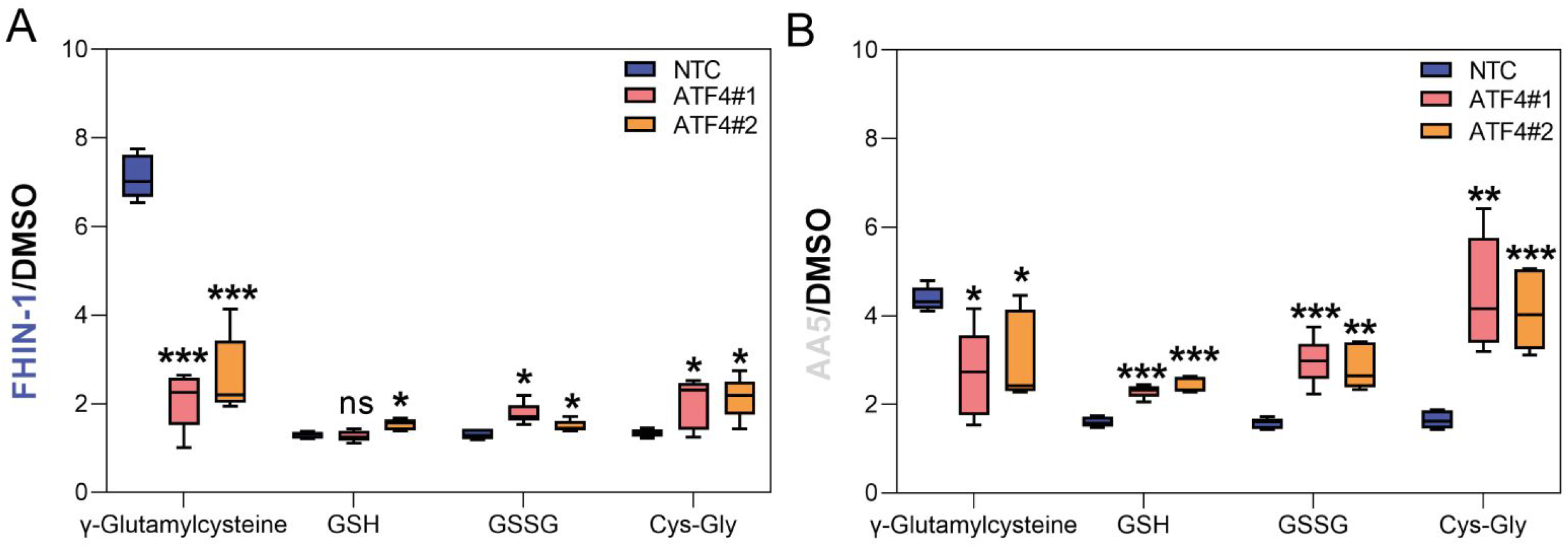
Supporting data for figure 7. (A) Interleaved box and whiskers plot of fold change in glutathione-related metabolites in DMSO-versus FHIN-1 (20 μM)-treated *Fh1^fl/fl^* cells with or without siRNA-mediated silencing of *Atf4* (24 h timepoint) (*n* = 5 independent biological replicates). (B) Interleaved box and whiskers plot of fold change in glutathione-related metabolites in DMSO-versus AA5 (1 μM)-treated *Fh1^fl/fl^* cells with or without siRNA-mediated silencing of *Atf4* (24 h timepoint) (*n* = 5 independent biological replicates). (A-B) p value determined by multiple unpaired t-tests, corrected with two-stage step-up method of Benjamini, Krieger and Yekutieli - FDR = 5%. p <0.05*; p <0.01**; p < 0.001***.

